# Single Cell Expression Analysis of Ductal Carcinoma in Situ Identifies Complex Genotypic-Phenotypic Relationships Altering Epithelial Composition

**DOI:** 10.1101/2023.10.10.561724

**Authors:** Xiaodi Qin, Siri H. Strand, Marissa R. Lee, Aashrith Saraswathibhatla, David G. P. van IJzendoorn, ChunFang Zhu, Sujay Vennam, Sushama Varma, Allison Hall, Rachel E. Factor, Lorraine King, Lunden Simpson, Xiaoke Luo, Graham A. Colditz, Shu Jiang, Ovijit Chaudhuri, E. Shelley Hwang, Jeffrey R. Marks, Kouros Owzar, Robert B. West

## Abstract

To identify mechanisms underlying the growth of ductal carcinoma in situ (DCIS) and properties that lead to progression to invasive cancer, we performed single-cell RNA-sequencing (scRNA-seq) on DCIS lesions and matched synchronous normal breast tissue. Using inferred copy number variations (CNV), we identified neoplastic epithelial cells from the clinical specimens which contained a mixture of DCIS and normal ducts. Phylogenetic analysis based on the CNVs demonstrated intratumoral clonal heterogeneity was associated with significant gene expression differences. We also classified epithelial cells into mammary cell states and found that individual genetic clones contained a mixture of cell states suggesting an ongoing pattern of differentiation after neoplastic transformation. Cell state proportions were significantly different based on estrogen receptor (ER) expression with ER-DCIS more closely resembling the distribution in the normal breast, particularly with respect to cells with basal characteristics. Using deconvolution from bulk RNA-seq in archival DCIS specimens, we show that specific alterations in cell state proportions are associated with progression to invasive cancer. Loss of an intact basement membrane (BM) is the functional definition of invasive breast cancer (IBC) and scRNA-seq data demonstrated that ongoing transcription of key BM genes occurs in specific subsets of epithelial cell states. Examining BM in archival microinvasive breast cancers and an *in vitro* model of invasion, we found that passive loss of BM gene expression due to cell state proportion alterations is associated with loss of the structural integrity of the duct leading to an invasive phenotype. Our analyses provide detailed insight into DCIS biology.

**SIGNIFICANCE:** Single cell analysis reveals that preinvasive breast cancer is comprised of multiple genetic clones and there is substantial phenotypic diversity both within and between these clones. Ductal carcinoma in situ (DCIS) of the breast is a non-invasive condition commonly identified through mammographic screening. A primary diagnosis of DCIS carries little mortality risk on its own, but its presence is a risk factor for subsequent clonally related invasive breast cancer (IBC) (1–5).

---

DCIS is a neoplastic proliferation of mammary epithelial cells within an intact basement membrane (BM). These lesions share virtually all of the genomic alterations found in IBC and therefore no specific genetic events have been associated with disease progression (6), although there are expression signatures that predict progression and benefit of radiotherapy (7,8). While mechanisms of invasion are not well understood, prior genomic analysis at the single epithelial cell level indicates that invasion is commonly polyclonal (9). This is consistent with no single event being responsible for the acquisition of the invasive phenotype. Processes that have been proposed as mechanisms of invasion include active degradation of the BM by the neoplastic cells, loss of myoepithelial cells which may serve as a cellular barrier to invasion, and mechanical pressures from the proliferative and expanding duct (10–14).

Single-cell RNA sequencing (scRNA-seq) has emerged as a valuable tool to understand tissue composition and expression at single-cell and whole-transcriptome resolution. A recent application of this approach to DCIS concluded that significant intra- and inter-tumoral heterogeneity exists (15); however, insights into the genesis of DCIS heterogeneity was limited by the lack of patient-matched normal samples. As part of the Human Tumor Atlas Network (HTAN), we focused on changes in cellular composition and phenotypes that occur between normal breast and DCIS starting from scRNA-seq data generated on DCIS and matched synchronous normal breast tissue. We observed significant alterations in epithelial cell states and transcriptome heterogeneity which accompanied intratumoral clonality. We extended the study to bulk RNA-seq data sets and analyses of DCIS and risk of progression to IBC in both patient data and an *in vitro* model. We hypothesized that altered composition of the DCIS ducts leads to a loss of ductal integrity and promotes subsequent invasion.

## RESULTS

### scRNA-seq distinguishes normal from neoplastic epithelial cells

We performed scRNA-seq on DCIS (n = 16) and matched synchronous normal tissue (n = 12) from 15 patients undergoing mastectomy (sample set 1). We filtered out 3 DCIS and 5 Normal samples based on QC metrics. The QC results for all samples are shown in **Supplementary Figure 1**, and **Supplementary Table 1** lists the 13 DCIS and 7 Normal samples from 14 patients that passed QC with patient demographic features. Both the DCIS and normal specimens were bisected, with one half used for single-cell dissociation and scRNA-seq, and the other half formalin fixed and embedded in paraffin (FFPE) to confirm the presence and extent of DCIS and normal breast epithelium (**Figure 1A**). From the FFPE samples, we performed whole-genome sequencing (WGS) and bulk-RNA sequencing on microdissected areas of histologically confirmed DCIS.

**Figure 1.**
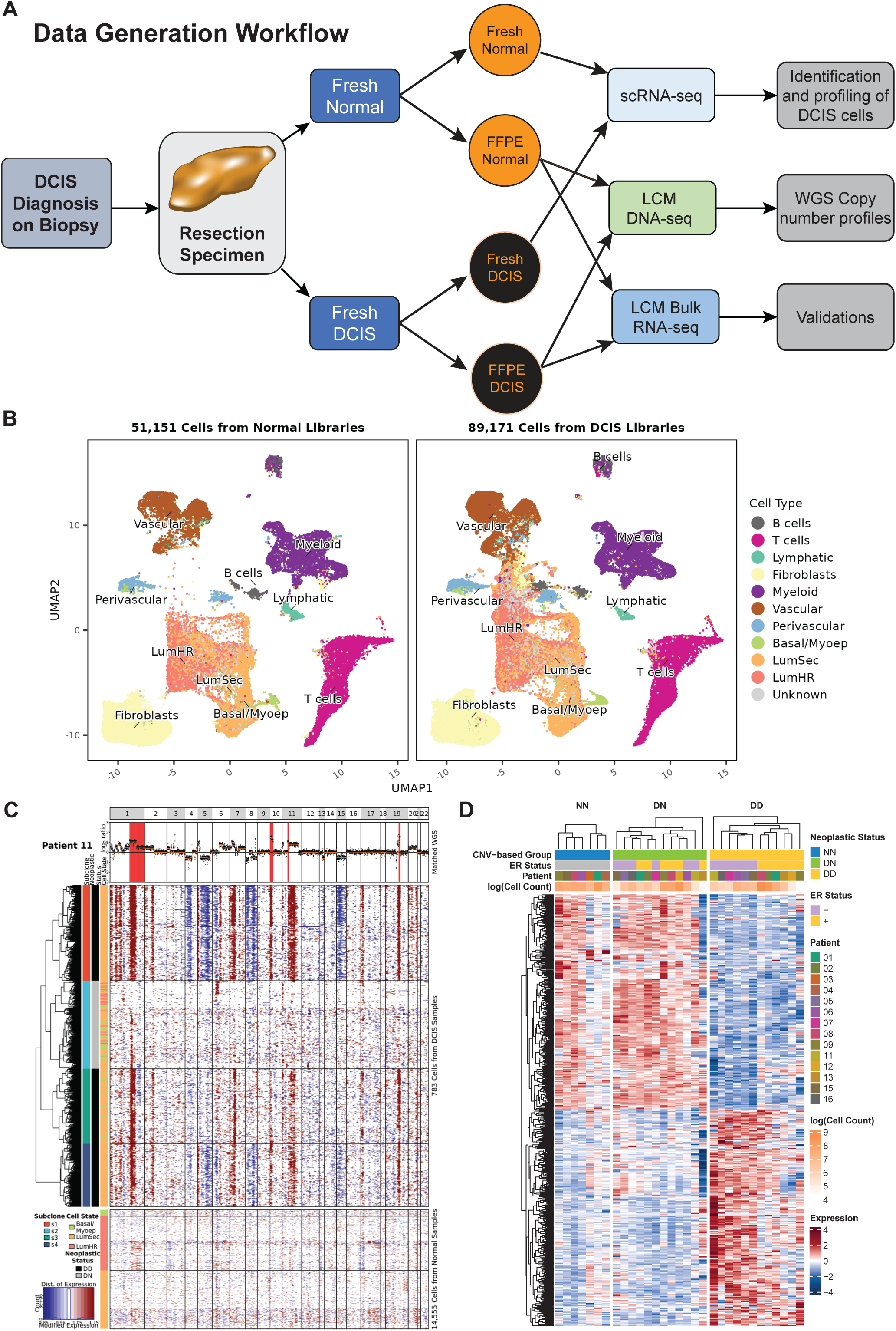
Overview of workflow. **A)** Laboratory workflow to obtaining multi-modal sequencing data from resected specimens (Sample set 1) to infer cell phenotypes from scRNA and DNA data, infer cell fractions from specimen-matched bulk RNA data for downstream analyses. **B)** UMAP embedding plots based on single-cell transcriptome profiles from 140,322 cells across 13 DCIS sample libraries (89,171 cells) and 7 normal breast sample libraries (51,151 cells) obtained from 14 patients. Data integration and dimension reduction was performed on the entire collection of cells to identify 11 primary cell types. After performing the embedding analysis on the latter, the cells were split depending on whether they were derived from a normal breast sample or DCIS sample library. The inferred cell type was used to color each coordinate. For each cell type, the ID is labeled at the median position of all cells in that category. **C)** A representative example of patient tumor-matched copy number motifs inferred from a low-pass WGS (upper panel) and inferred from single-cell DCIS (middle panel) libraries. A panel of 7 normal libraries were used as the reference (bottom panel). **D)** Relative expression of pseudo-bulk samples of epithelial cells from DCIS libraries inferred to be either DD (17,044 cells in 12 pseudo-bulk samples) or DN (14,309 cells from 12 pseudo-bulk samples), and cells from normal samples (NN, 14,555 cells in 7 pseudo-bulk samples). The top 250 up-regulated and 250 down-regulated differentially expressed genes (from DESeq2) ranked by adjusted *p*-value between DD and NN are displayed with expression of DN pseudo-bulk samples shown for comparison. Color bars indicate neoplastic status (DD, DN, NN), ER status (DD and DN samples only), patient ID, and cell count (natural log scale).

Two-dimensional UMAP embedding plots of cells from DCIS and matched normal tissue samples colored by cell type (**Figure 1B**) show that the overall architecture of the clustering was preserved between the two sample types, but that some clusters were only present in DCIS samples from specific patients (**Supplementary Figure 2A**). Based on histologic examination of the facing FFPE section, we recognized that the dissociated cells from DCIS samples were comprised of a mixture of neoplastic and non-neoplastic epithelial cells (**Supplementary Figure 2B**). We used the scRNA expression to infer cell-specific genomic copy number variation (CNV) profiles and compared those to WGS-based CNV profiles derived from microdissected DCIS from the adjacent paraffin block (**Figure 1C, Supplementary Figure 2C**). Epithelial cells with CNVs from DCIS specimens were categorized as DCIS cells (“DD” <u>D</u>CIS samples, <u>D</u>CIS cells) and those lacking these CNVs were categorized as “DN” (<u>D</u>CIS samples, <u>N</u>ormal cells) (decision rules for this categorization are contained in **Supplementary Table 2**). Additionally, since our study included a series of normal or uninvolved specimens, these epithelial cells were categorized as “NN” (<u>N</u>ormal samples, <u>N</u>ormal cells). To further validate that our inferred CNV-based approach distinguished DD from DN epithelial cells we compared aggregated gene expression (pseudo-bulk samples) between NN and DD cell populations. A heatmap of the top 250 significantly up-regulated and 250 down-regulated genes comparing the DCIS (DD) to normal cells (NN) is shown in **Figure 1D** (the complete list of differentially expressed genes is provided in **Supplementary Table 3**). DN cells were not used in the differential expression (DE) analysis for this heatmap but cluster closely with cells from normal breast (NN) (**Figure 1D**), indicating that the DN cells are admixed normal breast epithelial cells known to exist in these specimens based on histologic examination of the facing block. Comparison of DD to either DN or NN revealed similar pattern in significant pathway alterations, including cell cycle progression and metabolic pathways elevated in DD, further supporting the accuracy of the cell level categorization based on inferred copy number (**Supplementary Figure 2D**).

### Inferred CNV subclones demonstrate intratumoral genetic and expression heterogeneity

CNV analysis revealed that most of the DCIS cases contained cells with differing copy number states indicative of separate clones. The polyclonal nature of DCIS has been established (9) and our single cell analyses allowed us to explore the relationship of these clones with their phenotypic properties, including epithelial cell state. We first sought to identify these epithelial cell states using expression signatures based on the recent compendium of single cell data (16) from over 200 normal breast samples classifying the epithelium into three primary epithelial “states” (Basal, Luminal secretory (LumSec), and Luminal hormone responsive (LumHR)) and 11 “substates” of breast epithelium (1 Basal substate, 7 LumSec substates, and 3 LumHR substates). Based on the expression of characteristic markers, myoepithelial cells comprise the majority of the Basal cell state and substate, and are hereafter referred to as Basal/myoepithelial (Basal/Myoep) cells. From our scRNA data, two sample phylogenetic trees with cell states are shown in **Figure 2A** (other cases are shown in **Supplementary Figure 3A**). Sample 01D is an ER+ DCIS that demonstrates four subclones or branches of DCIS cells (and 1 branch of non-neoplastic DN cells) and Sample 15D is an ER-DCIS with three neoplastic clones. Fractional breast cell state composition of each of these branches indicates that none of these subclones are a monolithic population at the phenotypic level but that each subclone contains varying proportions of cell states and substates.

**Figure 2.**
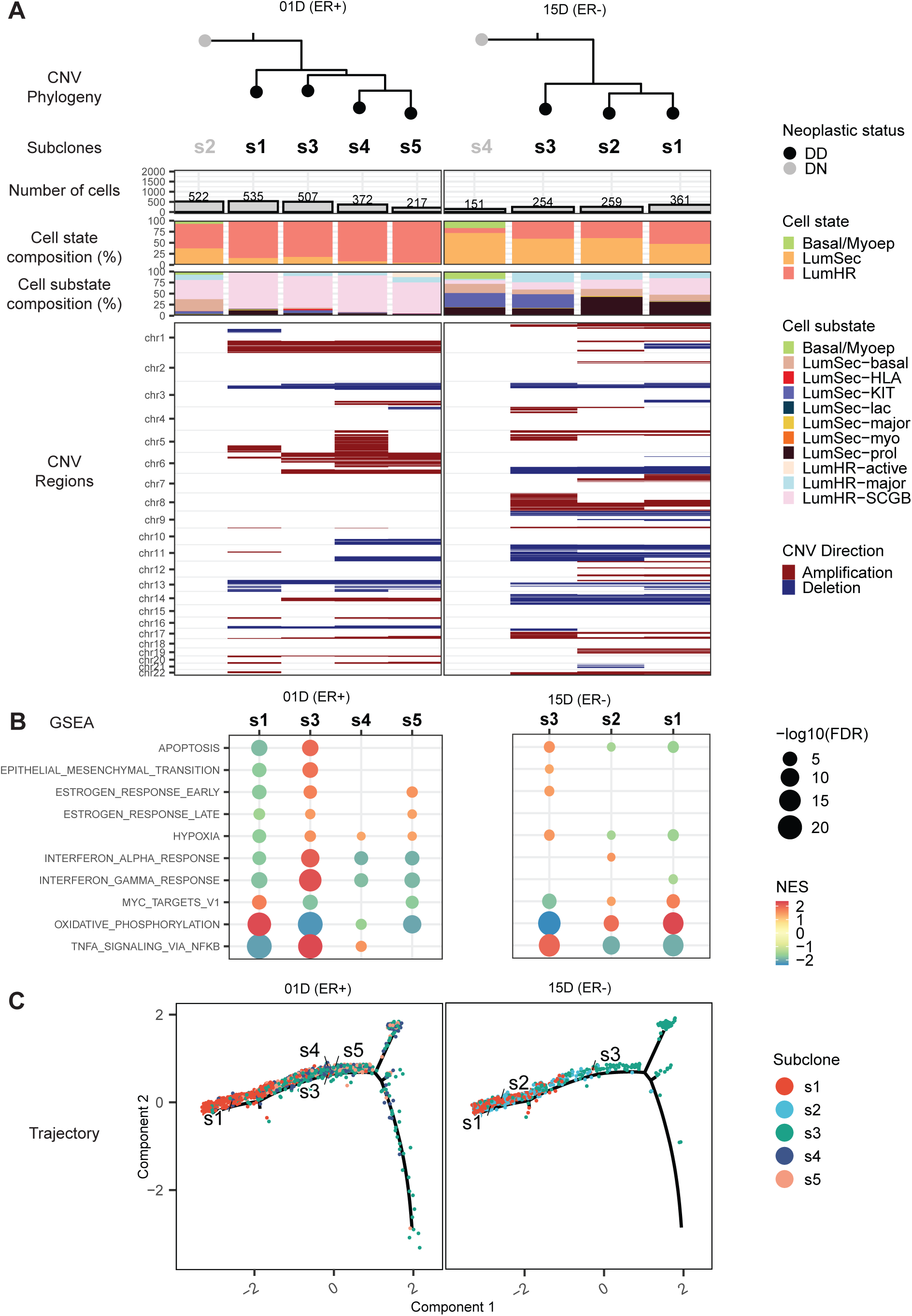
Heterogeneity of CNV profiles in DCIS. **A)** Phylogenetic trees constructed using inferred subclone CNVs show the relationship between subclones that were assigned neoplastic status classifications (DD versus DN) as shown by tree tips and subclone label colors. The number of cells, cell type compositions (states and substates), and inferred CNV regions by subclone are shown in the bar charts and tile plots. **B)** GSEA results for a given DCIS subclone as compared to others in the same sample are shown. **C)** Subclone states overlayed onto trajectory maps.

In considering the subclones as single populations, each subclone exhibited enrichment of unique combinations of Cancer Hallmark pathways (**Figure 2B** and **Supplementary Figure 3B**). Subclones tend to occupy specific positions on the composite trajectory map (**Figure 2C** and **Supplementary Figure 3C**) and the composite epithelial UMAP (**Supplementary Figure 3D**). In this way, we can visualize genotype-phenotype relationships at the level of cell state and specific gene expression pathways. From these analyses we concluded that subclones demonstrate unique variations in cell state heterogeneity and distribution. Further, between subclones, we observed highly significant pathway differences that are not readily evident either by examining cell state or UMAP position. For example, subclones of 01D exhibit similar cell state distributions but subclones S1 and S3 have several pathways that distinguish these two populations. Examination of individual cases reveals the complex genetic and phenotypic heterogeneity and indicates that clonal populations are themselves diverse mixtures of different mammary cell states.

### ER status distinguishes DCIS cell state composition

We next analyzed our cases aggregated by patient, neoplastic status, and estrogen receptor (ER) status. From among 140,322 cells which passed final quality control metrics for scRNA, we classified 45,908 epithelial cells from DCIS and normal libraries into three primary states and 11 substates as described above (16) (**Supplementary Figure 4A**). The three primary cell states are displayed in epithelial-specific UMAPs separated into DCIS (DD) and non-DCIS cells (DN/NN) (**Figure 3A**). We observed good concordance between the positions of the cells on the UMAP and the three cell states assigned independently based on sets of signature genes (16). Comparing DCIS (DD) to normal (DN/NN), we observed an overall loss of Basal/Myoep cells and LumSec in the DCIS cells compared to the normal cell population.

**Figure 3.**
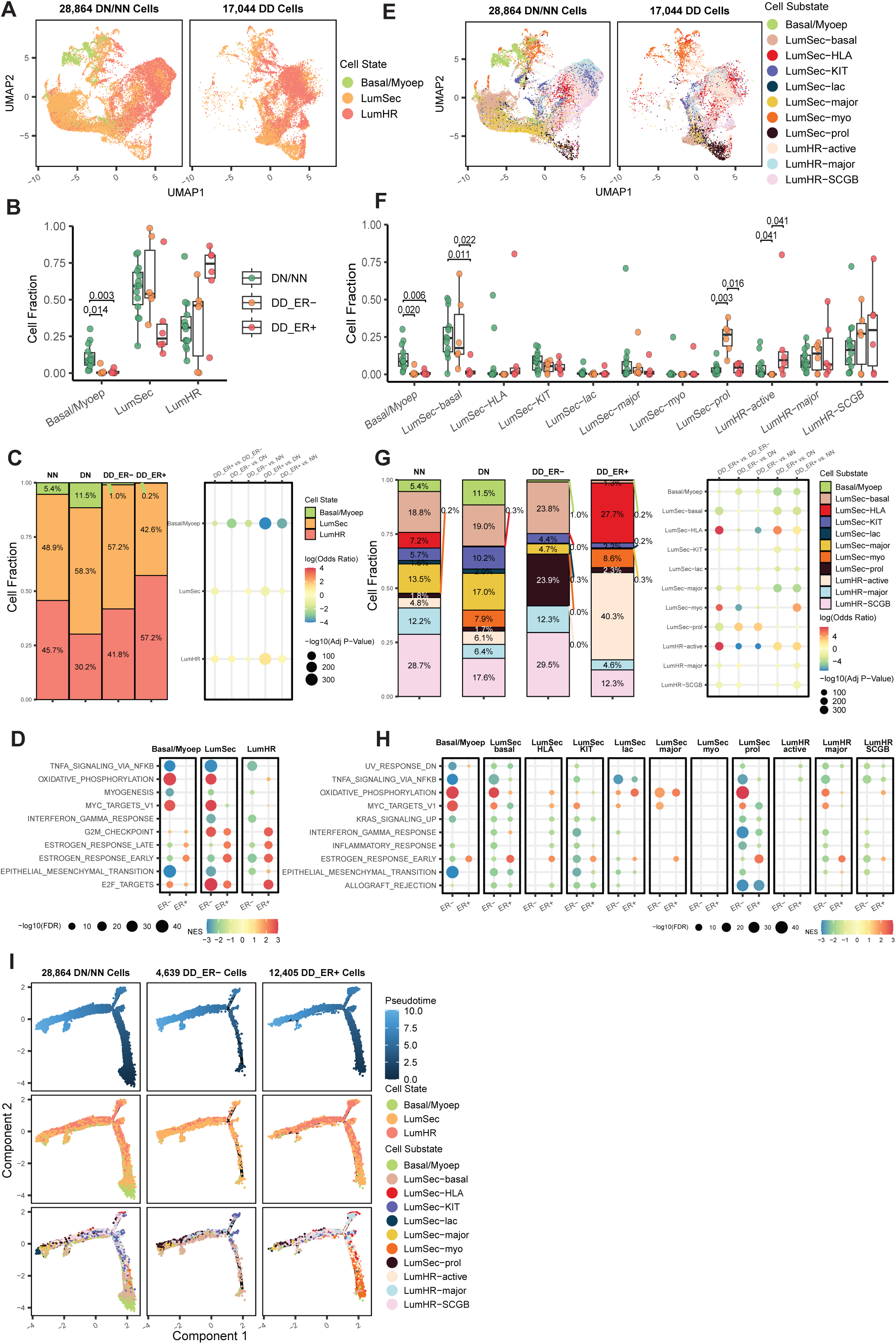
Identification and comparison of epithelial subtypes. **A)** UMAP embedding plots based on 45,908 epithelial cells (28,864 DNNN (left), 17,044 DD (right)) colored by the 3 major epithelial cell states found in normal breast. Data integration and dimension reduction was performed on epithelial cells prior to UMAP embedding. **B)** Relative cell fractions of 3 major epithelial cell states at the pseudo-bulk level for each sample for DN/NN, DD ER- and DD ER+. Variation in cell fraction by group was tested using Wilcoxon rank-sum and *p*-values were adjusted using Benjamini-Hochberg method to account for multiple comparisons. Results with an adjusted *p*-value of less than 0.05 are shown. **C)** Corresponding relative frequencies for the cell state compositions within DN, NN, DD ER+, and DD ER-groups are summarized as stacked bars with text showing the percentage of cells. The dot grid shows the resulting odds of cell state membership based on CNV and ER status. The color of the dots indicates the log odds ratio and the size indicates significance. **D)** Selected significant pathways from GSEA differentially enriched between pseudo-bulk samples from DN/NN and DD of 3 major cell states in ER-DCIS and ER+ DCIS, analyzed separately. The color of the dots indicates the normalized enrichment score and the size indicates significance measured as the negative log10 adjusted *p*-value. Panels **E**, **F**, **G** and **H** present results at finer resolution, where 11 epithelial cell substates are considered instead of the 3 major states. They correspond to panels **A**, **B**, **C** and **D**, respectively. **I)** Cell trajectory analysis based on DN/NN, DD ER- and DD ER+. Cells in the top row are colored by pseudotime, in the middle row cells are colored by the 3 major epithelial cell states, and in the bottom row cells are colored by the 11 epithelial cell substates.

We analyzed this further at the patient level and confirmed both a lower proportion and lower absolute number of Basal/Myoep cells in both ER+ and ER-DCIS compared to normal samples (**Figure 3B**, adj. *p* = 0.003, 0.014, respectively). LumHR cells comprised the greatest proportion of the DCIS epithelial cell population from ER+ DCIS and, concomitantly, there were fewer LumSec and Basal/Myoep cells in the DD populations. Interestingly, the ER-DCIS cells exhibited higher proportions of LumSec fractions compared to the ER+ DCIS. Aggregating the cell data confirmed highly significant distributional differences between normal (NN or DN), ER+ DCIS, and ER-DCIS (**Figure 3C**). Pathway analysis revealed substantial differences between the neoplastic (DD) and normal (DN/NN) in both ER+ and ER-DCIS (**Figure 3D, Supplementary Figure 4B, Supplementary Table 4)**.

We next repeated this analysis by applying the more granular 11 epithelial substates (UMAP in **Figure 3E**). Comparisons between normal, ER+ DCIS, and ER-DCIS cell populations demonstrated significant differences in cell substate compositions and expression pathways. (**Figure 3F-H**). The relative proportions of both the LumSec-basal and LumSec-prolif substates were higher in ER-DCIS compared to ER+ DCIS (adj. *p* = 0.022, 0.016, respectively). With respect to the LumHR substates, the relative proportion of the LumHR-active substate was significantly lower in ER-compared to ER+ DCIS and normal (adj. *p* = 0.041, 0.041, respectively). ER-DCIS contained substantial proportions of the various luminal hormone responsive subtypes (i.e., LumHR-major and LumHR-SCGB). Therefore, even though the ER-DCIS cells do not express appreciable levels of the estrogen receptor, a proportion of these cells retained expression profiles that classified them as LumHR.

Pathway analysis comparing ER+ and ER-DCIS to normal demonstrated global differences in TNF signaling (lower in ER-DCIS) and oxidative phosphorylation (higher in ER+ DCIS) across multiple cell substates (**Figure 3H, Supplementary Figure 4C, Supplementary Table 5).** We also noted some significant substate-specific differences between ER+ and ER-DCIS, compared to normal, including lower expression of the Epithelial Mesenchymal Transition pathway in the basal substate as well as lower Interferon Gamma Response in the LumSec-prolif substate in ER-DCIS compared to ER+ DCIS.

Next, we analyzed the progression of the differentiated states of the epithelial cells using a trajectory analysis (**Figure 3I**). The full branching pattern was established using all epithelial cells from DCIS and normal samples and then divided into three categories, normal (DN/NN), ER-DCIS and ER+ DCIS for the 3 cell states (middle panels) and 11 substates (bottom panels). Most Basal/Myoep cells occupied a branch at the earliest pseudotime whereas luminal cells (both LumSec and LumHR) were located throughout the timeline. Surprisingly, the ER-DCIS demonstrated a more restricted distribution with diminished number of cells at the early pseudotime branch compared to ER+ DCIS and normal epithelium implying that there is a differential accumulation of specific tumor cell populations over time.

### Cell state analysis in independent DCIS cohorts indicates outcome differences

To extend these findings, we used deconvolution to estimate the cell state composition from two additional data sets (sample set 2 and 3) of bulk RNA-seq derived from archival specimens. **Sample set 2** is RNA-seq from laser capture microdissected (LCM) epithelium of synchronous samples of normal, DCIS, and IBC from the same patient. **Sample set 3** is RNA-seq from macrodissected specimens from two previously reported longitudinal cohorts of DCIS with known disease outcomes (7) (**Supplementary Table 6**). We first confirmed the validity of the deconvolution method by comparing cell composition from bulk RNA-seq (**Figure 4A**) to the matched scRNA-seq results (**Figure 3B**). Correlating with the findings from scRNA-seq, we found lower relative proportion of Basal/Myoep cells in both ER+ and ER-DCIS vs normal samples (adj. *p* = 0.028, adj. *p* = 0.028) and differences between ER- and ER+ DCIS in LumSec (adj. *p* = 0.022) and LumHR (adj. *p* = 0.022). At the individual sample level, we observed significant correlation between scRNA-seq and FFPE bulk RNA-seq data for all three epithelial cell states (**Supplementary Figure 5A**).

**Figure 4.**
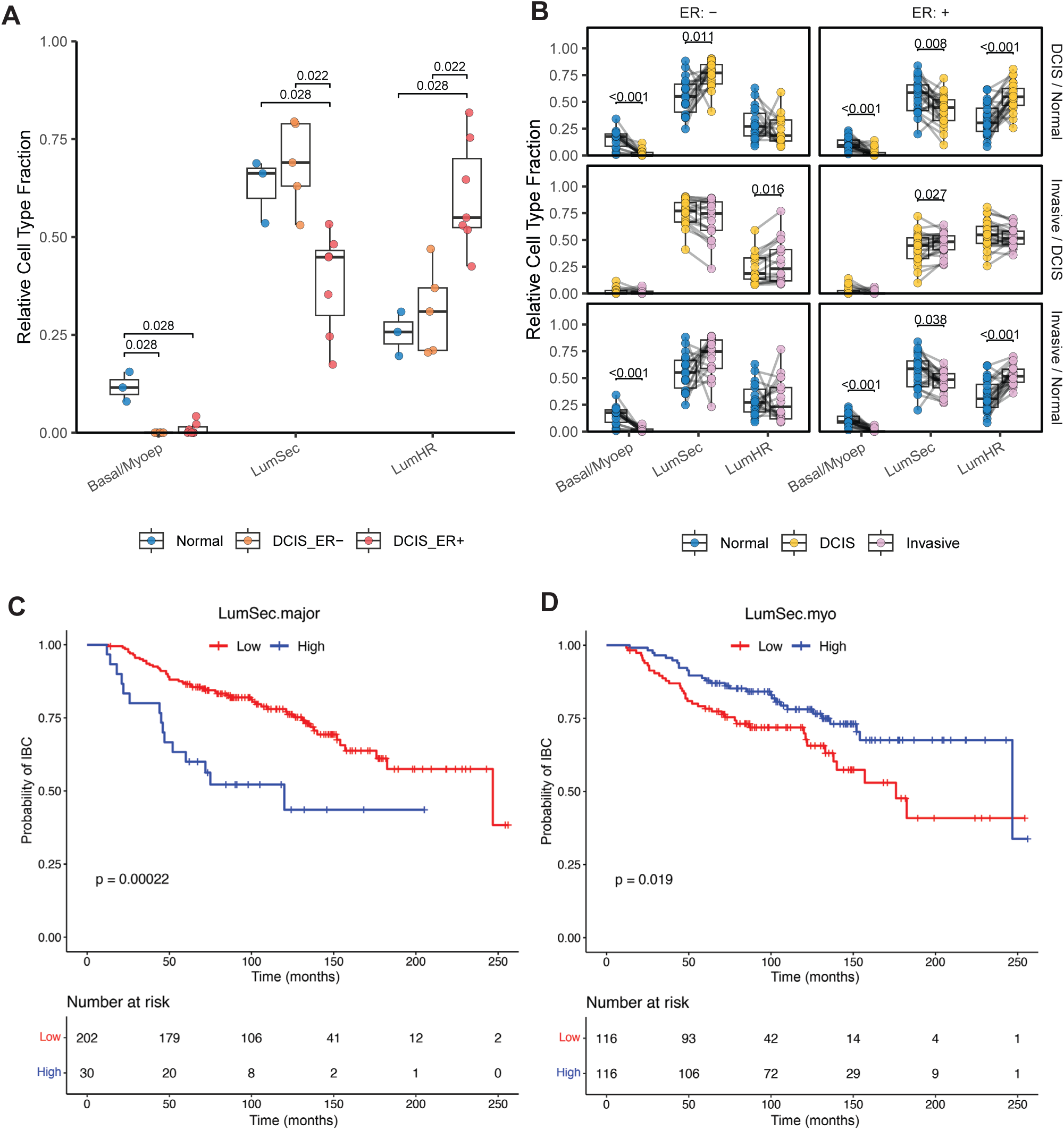
Analysis of cell type proportions in independent DCIS cohorts. **A)** Analysis of imputed cell fractions based on 3 major epithelial cell states found in normal breast from bulk-RNA sequencing datasets of the FFPE sample set matching the scRNA-seq samples as depicted in Figure 1A with 3 Normal, 7 ER+ DCIS, and 5 ER-DCIS libraries obtained from 11 patients (Sample set 1). Differences were tested using the unpaired two-sample Wilcoxon rank-sum test. *P*-values were adjusted using Benjamini-Hochberg method to account for 9 comparisons. Results with an adjusted *p*-value of less than 0.05 are shown. **B)** Analysis of imputed cell fractions based on 3 major epithelial cell types found in normal breast from bulk-RNA sequencing datasets of 126 patient-matched normal breast, DCIS, and IBC libraries (24 ER+ DCIS, 18 ER-DCIS, 42 matched normal, and 42 matched IBC) (Sample set 2). Points for each patient are connected with a grey line. Pairwise differences were tested using one-sample Wilcoxon signed-rank tests. *P*-values were adjusted using Benjamini-Hochberg method to account for 18 comparisons. **C, D)** Kaplan-Meier curves of time to IBC recurrence for high and low levels of LumSec-major (**C**) and LumSec-myo (**D**), from the analysis of imputed cell fractions bulk-RNA sequencing datasets of retrospective DCIS case – control cohorts (Sample set 3). The *p*-values were derived from Cox proportional hazard models to assess the difference in recurrence risk between the two groups. The table below each plot shows the number of patients still at risk of recurrence at each time point.

In sample set 2 (synchronous), the relative proportion of Basal/Myoep cells was again significantly lower in DCIS vs normal breast for both ER+ and ER-DCIS (adj. *p* < 0.001, **Figure 4B**). In addition, LumHR cells were significantly enriched in ER+ DCIS compared to the matched normal samples (adj. *p* < 0.001), but this was not observed for ER-DCIS. Conversely LumSec cells were significantly less abundant in ER+ DCIS compared to matched normal breast (adj. *p* = 0.008) but increased in ER-DCIS (adj. *p* = 0.011). Comparing the matched and synchronous DCIS to IBC, we observed minor increases of LumSec in ER+ and LumHR in ER-IBC. Overall, differences in cell state distribution between synchronous DCIS and IBC are much less pronounced than between normal and DCIS.

To investigate the association between the relative abundance of the cell states and substates and risk of invasive progression in DCIS, we analyzed the imputed cell fractions in retrospective DCIS case–control cohorts of patients with a primary diagnosis of DCIS who later either did or did not have IBC progression (sample set 3, longitudinal follow up). We found that patients with high levels of LumSec-major cells (*p* = 0.0002), and low levels of LumSec-myo cells (*p* = 0.019) were associated with shorter time to IBC recurrence (**Figure 4C, 4D, and Supplementary Figure 5B**). Other cell states and substates were not significantly associated with invasive progression.

### Basement membrane gene expression in DCIS

BM maintenance in the breast is an ongoing and active process (17) and a loss of BM leads to failure of developmental epithelial structures (18–20). Experimental studies have shown that destruction of the BM is associated with genetic instability and mammary tumorigenesis (21). Thus, we next evaluated whether differences in epithelial state composition could influence the production and integrity of the BM. In our scRNA data, we examined the expression of canonical BM genes (GO:0005604) (22) within the epithelial cell states, and fibroblasts (**Figure 5A**). As expected, fibroblasts demonstrated relatively high expression of many of these genes. Notably, Basal/Myoep cells also demonstrated significant expression of a set of BM genes, indicating that this epithelial cell state contributes to the ongoing synthesis of the BM. While there was overlap, expression of some BM genes was uniquely enriched in each cell state, such as *LAMC2* in Basal/Myoep cells. Among the 11 substates, several had prominent BM gene expression including LumSec-basal and LumSec-KIT (**Supplementary Figure 6A**).

**Figure 5.**
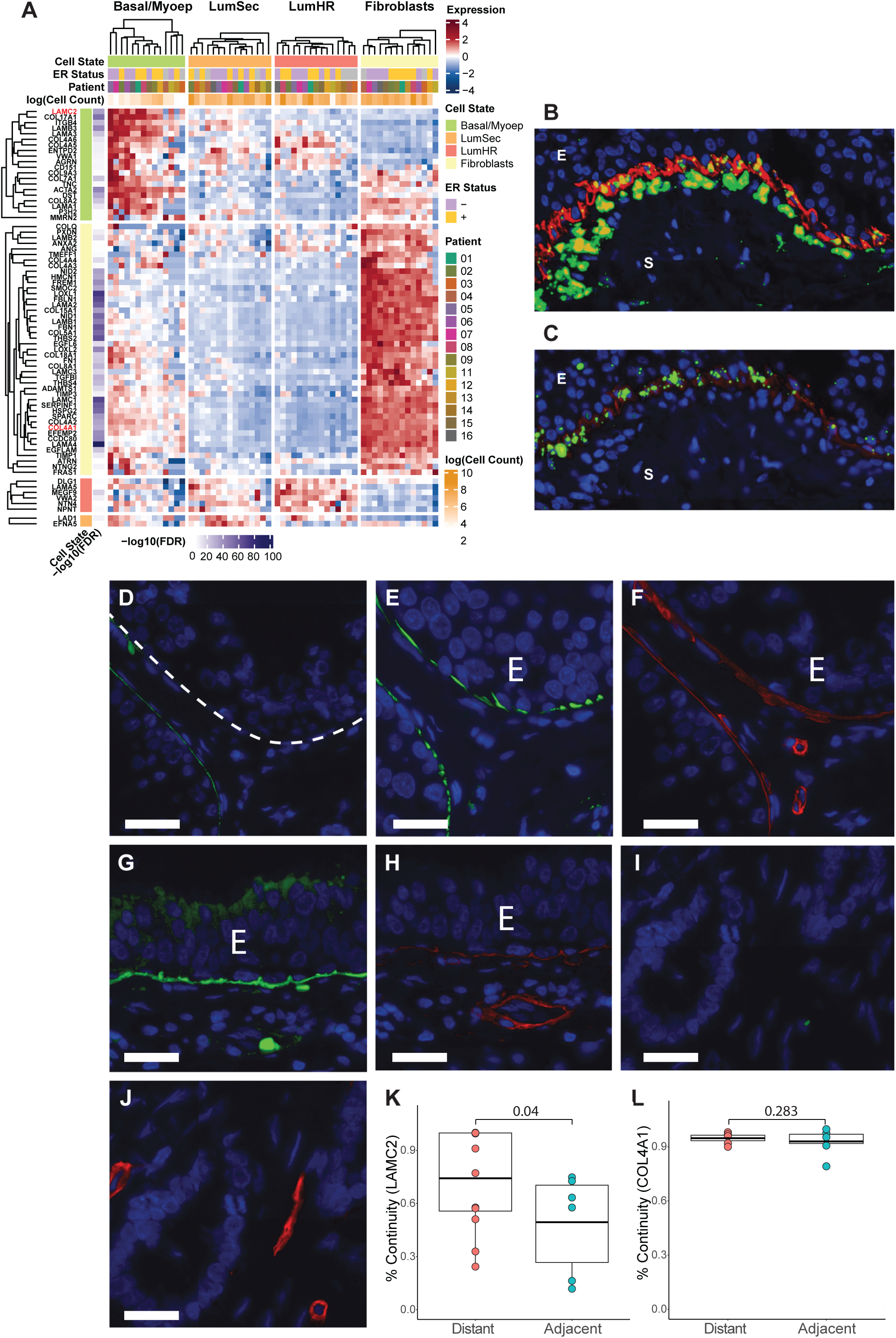
Basement membrane expression in DCIS. **A)** Relative expression of BM genes across the three major cell states and fibroblasts within each pseudo-bulk samples from scRNA-seq data (Sample set 1). Displayed genes were most significantly up-regulated (adj. p < 0.01) in each specific cell type compared to all other cell types. **B)** Detection of COL4A1 RNA (green) in DCIS epithelium (E) and adjacent stroma (S) with ACTA2 protein (red) expression demarcating the basal zone of the DCIS sample. **C)** Detection of LAMC2 RNA (green) in DCIS epithelium (E) and adjacent stroma (S) with ACTA2 protein (red) expression demarcating the basal zone of the DCIS sample. **D)** Immunofluorescence of LAMC2 in DCIS (E) located adjacent to invasion (Sample set 4). **E)** Immunofluorescence of ACTC2 in DCIS (E) located adjacent to microinvasion. **F)** Immunofluores-cence of COL4A1 in DCIS (E) located adjacent to invasion. **G)** Immunofluorescence of LAMC2 in DCIS (E) located distant to invasion. **H)** Immunofluorescence of COL4A1 in DCIS (E) located distant to invasion. **I)** Immunofluorescence of LAMC2 in IBC. **J)** Immunofluorescence of COL4A1 in IBC. **K)** Comparison of percentage of continuity for LAMC2 in DCIS ducts between DCIS adjacent to invasion or distant to invasion. **L)** Comparison of percentage of continuity for COL4A1 in DCIS ducts between DCIS adjacent to invasion or distant to invasion. Scale bar = 20 um.

*In situ* analysis of protein and RNA expression of selected BM genes provided additional evidence that Basal/Myoep cells contribute to the formation and maintenance of the BM (**Figure 5B and 5C**). *COL4A1* RNA expression was predominantly found in fibroblasts adjacent to the epithelial compartment (**Figure 5B**). Moreover, substantial expression of *COL4A1* was also evident in a subset of the ACTA2 (smooth muscle actin)-positive cells (i.e., Basal/Myoep cells) at the edge of the epithelial compartment. *LAMC2 RNA* was expressed in Basal/Myoep cells but not fibroblasts (**Figure 5C**). Based on these data, the epithelial compartment appears to play an active role in both the maintenance and loss of integrity of the BM.

We further investigated whether BM integrity is associated with breast cancer invasion by examining the physical continuity of specific BM components in a series of DCIS adjacent to areas of microinvasion (defined as ≤1 mm invasive component, sample set 4, n = 8). LAMC2 expression was greatly decreased and sometimes absent in some ducts adjacent to the invasion (**Figure 5D**) despite the presence of Basal/Myoep cells as defined by ACTA2 expression (**Figure 5E**). Conversely, COL4A1 protein was found in a consistent pattern around the ducts adjacent to the invasive component (**Figure 5F**). In contrast, DCIS distant from the invasive component had more continuous expression of LAMC2 and COL4A1 (**Figure 5G and 5H**). The BM distribution of COL4A1 and LAMC2 is lost in the invasive component of all samples (**Figure 5I and 5J**). Quantitative measurements of LAMC2 continuity demonstrated a significantly lower percentage of continuity comparing DCIS adjacent to IBC versus DCIS distant (*p* = 0.04) whereas COL4A1 did not show this difference (**Figure 5K and 5L**). Normal breast demonstrated a continuous COL4A1 and LAMC2 layer surrounding all identified ducts and lobules (**Supplementary Figure 6B and 6C**). This shows that DCIS can exist without an intact LAMC2 layer and loss of this BM component is associated with DCIS near microinvasion suggesting that this may be an early step in progression. COL4A1, the essential structural feature of the BM (19), is lost at the invasive step.

To test the role of BM integrity in regulating the invasive phenotype, we used our previously established three-dimensional *in vitro* culture model in which MCF10A mammary epithelial acini are encapsulated in stiff hydrogels (elastic modulus of 2.4 kPa) containing interpenetrating networks of reconstituted BM matrix and alginate (23). In this model, the increased stiffness relative to normal breast promotes an invasive phenotype in some of the acini. The acini, formed from single MCF10A cells, exhibit a continuous BM at the periphery including layers of laminin-332 and COL4 (**Figure 6A**) mimicking mammary duct BM. To perturb the integrity of the BM, the acini were treated with collagenase-IV for 1 hour degrading the COL4 protein (**Figure 6B; Methods**) and decreasing the thickness of the BM (**Figure 6C; Methods**). Control and collagenase-treated acini were then encapsulated in the interpenetrating hydrogels for 7 days. On day 7, we observed that the acini with the degraded COL4 layer were more invasive (**Figure 6D-E**) and showed decreased circularity (**Figure 6F; Methods**), indicating that loss of this key component of the BM promotes the invasive phenotype.

**Figure 6.**
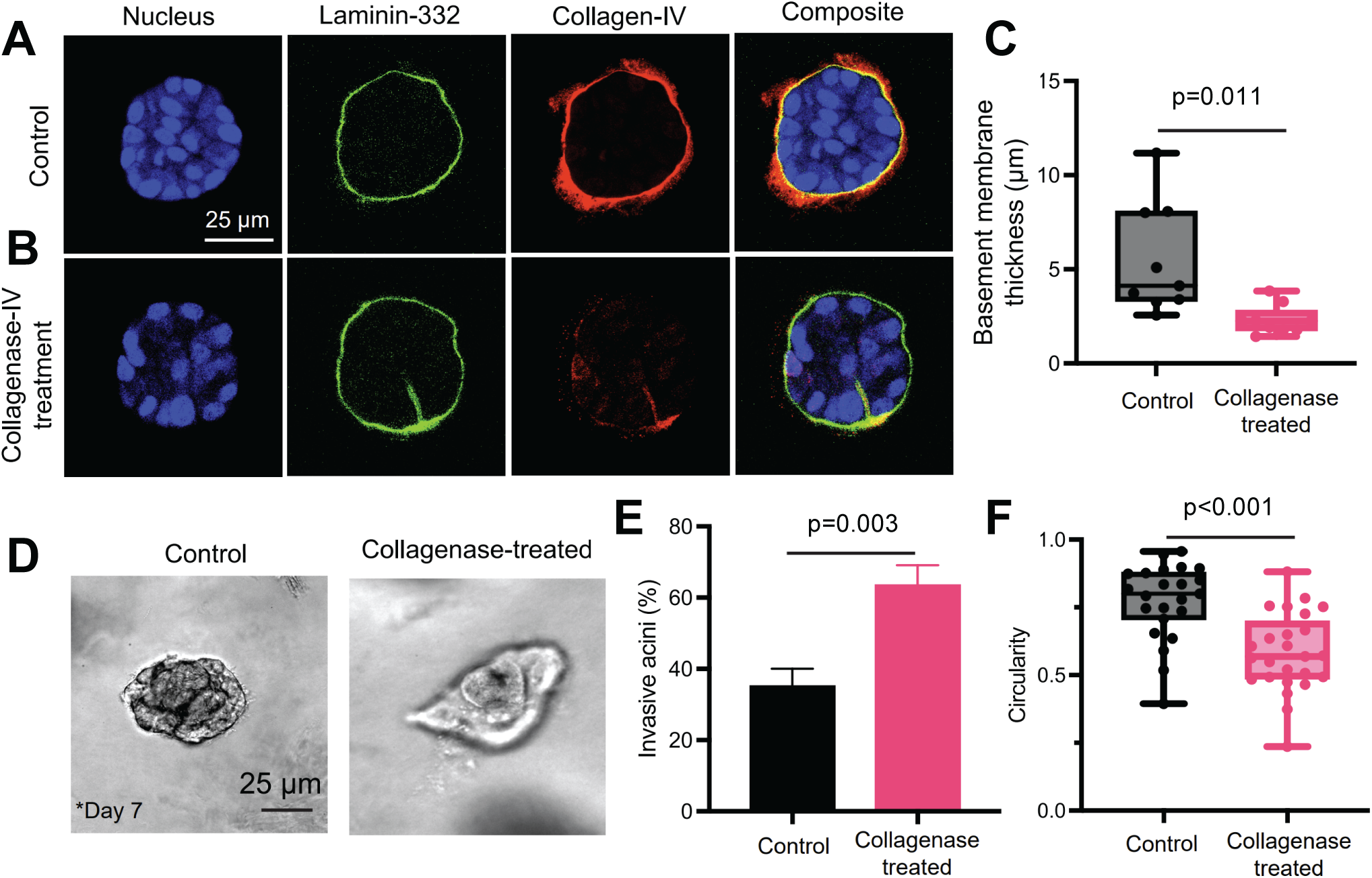
Compromising BM integrity by degrading collagen-IV increased the invasion of mammary epithelial cells in a 3D in vitro culture model. **A)** Confocal images of nuclei, laminin, and collagen-IV of control MCF10A acini. **B)** Confocal images of nuclei, laminin, and collagen-IV of collagenase-IV-treated acini. **C)** Thickness of BM in control and collagenase-treated conditions. Each dot corresponds to an acini (n ≥ 9 acini). **D)** Phase contrast images of acini on day 7 after encapsulation in control (top) and collagenase-treated (bottom) conditions. **E)** Percentage of invasive acini in control and collagenase-treated conditions (n ≥ 2 different gels). **F)** Circularity of acini in control and genase-treated conditions. Each dot corresponds to an acini (n ≥ 18 acini). Two-sided Welch’s t-test was performed for panels **C**, **E**, and **F**.

### A conceptual model for compromise of epithelial integrity

A conceptual model of cancer progression that incorporates the consequences of decreased Basal/Myoep cell proportions in DCIS is presented in **Figure 7**. We hypothesized that the relative expansion of neoplastic luminal cells leads to an alteration of the epithelial microenvironment. The relative reduction of cell states (Basal/Myoep and LumSec-basal) decreases the contributions made by the epithelium to the BM. This results in a decrease in the structural integrity of the DCIS-involved duct. This loss of integrity results in cells losing contact with the epithelium compartment and entering the stromal compartment. This passive, loss-of-function event is interpreted as invasion.

**Figure 7.**
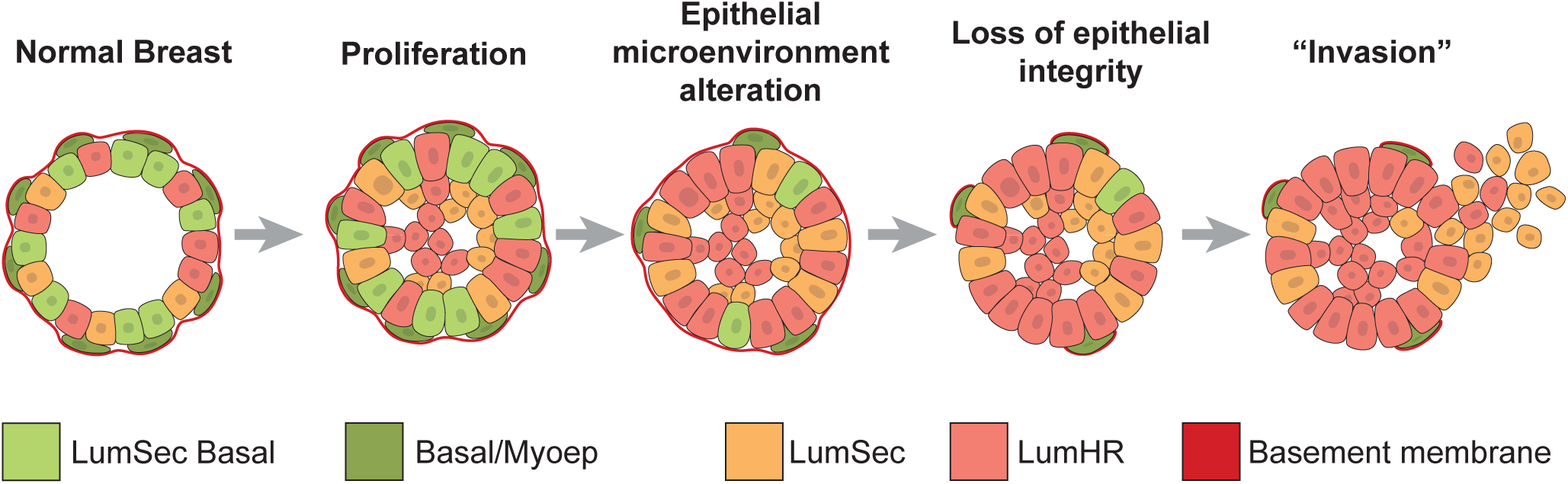
Conceptual model of DCIS invasion. Hypothesized changes in epithelial cell fractions during disease progression from normal breast to early and late DCIS stages.

## DISCUSSION

We show that DCIS is comprised of multiple subclones with significant gene expression differences. Further, the DCIS subclones are themselves not monolithic based on their composition of mammary cell states. Normal mammary ducts and lobules are composed of several spatially organized cell types that exist along a differentiation spectrum starting from pluripotent stem cells (24) and a defining feature of DCIS is that it recapitulates normal breast ductal growth patterns, strongly suggesting that the cells retain the ability to differentiate along similar lines. For this reason, in addition to identifying subclones, we also chose to classify the DCIS epithelial cells into phenotypic states according to the recently published and most extensive single cell atlas of normal human breast cells, comprising 200 normal breast tissues (16). This resource defines 3 states and 11 substates of epithelial cells that we used to show that genetic subclones of DCIS contain multiple cell states and that each cell state can be made up of multiple genetic clones.

A key feature of the current study is the discrimination of epithelial cells from the DCIS specimens as “contaminating” normal epithelial cells versus those that are part of the cancer. The use of inferred copy number based on gene expression is a validated approach for single cell data and our bulk whole genome sequencing from the same specimens confirms the presence of the CNV’s. Based on histology and inferred copy number, the sample in our study are comprised of highly varied mixtures of normal and DCIS indicating that analyses of these data without this step would be misleading.

We found that most of the DCIS lesions in the current study are composed of a mixture of multiple genetic clones that exist in a range of differentiation states. We observed notable differences in both the expression and relative abundance of cell states and substates between normal and DCIS cells, as well as between ER+ and ER-DCIS cells. Cells in the Basal/Myoep spectrum are significantly less abundant in the neoplastic cell populations, indicating that cell differentiation into this cell state is uncommon in DCIS. However, ER-DCIS (but not ER+) retained a significant proportion of cells with basal characteristics (LumSec-basal) distinct from the mature myoepithelial population. We concluded that each DCIS lesion has variable compositions of cell states that only in part mimic the distribution and heterogeneity of the normal breast, with ER-DCIS (compared to ER+ DCIS) more closely resembling the cell distribution in the normal breast. In a DCIS cohort with longitudinal outcome, we found that high levels of LumSec-major and low levels of LumSec-myo are associated with shorter time to IBC progression. LumSec-major cells express MMP7 which may promote invasiveness, whereas LumSec-myo cells share properties with mature myoepithelial cells and may retard progression (16).

Previous studies have found that attenuation of the Basal/Myoep cell layer is common in DCIS and has been considered as one of a number of potential mechanisms of invasion (12–14). However, while invasive cancers lose the Basal/Myoep layer, we recently showed, using spatial proteomics, that attenuation of the Basal/Myoep layer (defined by ACTA2 expression) is not associated with longitudinal progression of DCIS suggesting that there may not be a direct cause and effect relationship (25). Our current results show that the changes in cell composition related to the expansion of DCIS cells may affect normal epithelial function, in particular the structure and composition of the BM.

Some of the essential functions of the BM in maintaining integrity include providing a scaffold for the epithelial cells, helping them maintain their shape and resist mechanical stress, and anchoring the epithelial cells to the underlying matrix (18,26–28). In a mouse DCIS model, BM maintenance was shown to be an ongoing and active process (17). Using a 3D *in vitro* culture model, we demonstrated that compromising BM integrity by degrading COL4 in mammary acini increased the invasion of the mammary cells. COL4 is one of the main structural proteins in the BM and its importance in maintaining epithelial integrity is illustrated by its evolutionary appearance at the transition from unicellular to multicellular organisms (29). Notably in our investigation of BM continuity in DCIS adjacent to areas of microinvasion, we did not see COL4 loss in DCIS, only where there was invasion. These findings are consistent with previous studies on synchronous DCIS and IBC lesions which found epithelial gene expression changes indicative of progressive loss of basal layer integrity (30).

Our data support that overgrowth of neoplastic cell populations and the relative reduction of BM producing cells within the epithelium may create an unstable epithelial structure. This reframes the concept of invasion as a loss of function event due to an imbalance in critical cell populations essential for BM integrity during the neoplastic process, rather than a gain of function by the neoplastic cells. This model helps to explain the previously described genomic identity between epithelial cells in DCIS and invasive cancer and suggest novel alternative targets for future interception efforts that focus on maintenance of the BM.

## Methods

### Sample sets (Supplementary Table 6)

Sample set 1 (scRNA-seq set) consisted of DCIS (n = 16), IBC (n = 2) and synchronous normal tissues (n = 12) from 16 patients undergoing mastectomy at Duke and Stanford, collected between 2019 – 2023. Due to low sample size, the invasive samples were not included in these analyses. Both the DCIS and normal specimens were bisected with one half used for single-cell RNA sequencing, and the other half formalin fixed and embedded in paraffin (FFPE). Sections from the FFPE samples were used for histologic examination to confirm the presence and extent of DCIS and normal breast epithelium.

Sample set 2 (matched and synchronous breast specimens) consisted of FFPE tissue samples from 42 patients operated on at Stanford Hospital with matched and synchronous areas of normal breast, DCIS, and IBC.

Sample set 3 (DCIS with longitudinal outcome) consisted of 232 patients from the combined TBCRC and RAHBT cohorts included in the HTAN DCIS Atlas (7). These samples are the subset derived from patients with either no recurrence (n = 163) or IBC progression (n = 69).

Sample set 4 (DCIS with microinvasion) consisted of FFPE specimens with DCIS that exhibited areas of microinvasive cancer (<1mm) collected from 13 patients at Stanford Hospital.

### ScRNA-seq assay

Fresh tissue samples were collected within 1 h after devascularization and were immediately minced on ice before suspension in MACS Tissue Storage Solution (Miltenyi Cat# 130-100-008) in a 2 ml cryovial (Sarstedt Cat# 72.694.406) and stored at -80° C while awaiting pathological confirmation of DCIS in the matching DCIS FFPE H&E tissue section. Following confirmation, minced tissue samples were thawed in water bath and transferred to a gentleMACS C tube (Miltenyi Cat# 130-093-237) with enzymes from the Miltenyi Human Tumor Dissociation Kit, (130-095-929) according to protocol with gentleMACS Octo Dissociator with Heaters (130-096-427) program *37C_h_TDK_3*. Cells were washed using 3 mL Gibco RPMI 1640 Medium (ThermoFisher Scientific cat# 11875119), strained using MACS smart strainer 70 µm, and counted on a Countess II cell counter (ThermoFisher Scientific). Cells were resuspended in Hanks’ Balanced Salt Solution (ThermoFisher Scientific Cat# 88284) to 1200 cells/uL for immediate downstream processing. GEMs were generated and barcoded using the Chromium Single Cell 3ʹ _v3 reagents and workflow (10x Genomics) at Stanford Genome Sequencing Service Center as a commercial service. Libraries were sequenced on an Illumina NovaSeq.

### Low-pass whole-genome DNA-sequencing (WGS DNA-Seq) assay

Genomic DNA was isolated from FFPE samples using PicoPure DNA Extraction kit (Thermo Fisher Scientific # KIT0103). DNA library construction and sequencing were performed as previously published (7).

### Bulk RNA-seq

The bulk-RNA sequencing libraries were prepared from dissected areas of FFPE sections for sample sets 1, 2, and 3 as described previously (7) and then prepared for sequencing using the SMART-3SEQ protocol (31). Libraries were sequenced on an Illumina NextSeq 500 instrument with a High Output v2.5 reagent kit (Illumina # 20024906).

### Immunofluorescence (IF) on pathology specimens

IF staining was performed on paraffin-embedded tissue microarray (TMA) sections (4 µm) (Sample set 4). Briefly, slides were deparaffinized and hydrated using a xylene-ethanol series. Antigen retrieval was carried out at 116°C for 3 min. in a decloaking chamber in Antigen Unmasking Solution, Citrate-based pH6, at 10mM (Cat No. S236984-2 Agilent, Santa Clara, CA) for Col IV (Sigma Cat# AB769; 1:50) and Acta2 (ThermoFisher Cat# 53-9760-82; 1:3000) detection. For Lamc2 (SantaCruz Biotech Cat# sc-28330; 1:50), proteinase K (Agilent Tech Cat# S302080-2) digestion was done. Slides were incubated with primary antibody at room temperature for 45 min. For detection, Alexa Fluor conjugated secondary antibodies at 1:700 dilution were used. Counter staining was performed using Prolong Gold Antifade with DAPI (ThermoFisher Cat# P36935). Slides were scanned with a Leica Microscope slide scanner using Ariol Software (Leica Biosystems, CA).

### Combined immunofluorescence and RNA in situ hybridization assay

Co-detection of ActA2 protein and COL4A1 and LAMC2 RNA were performed using all ACD guidelines for performing in situ hybridization with the RNAscope 2.5 HD Detection Kit-RED (Cat# 322360) and RNA-Protein Co-Detection Ancillary kit (Cat# 323180) immunochemistry on FFPE TMA tissue sections (Sample set 4). The following RNA probes were used: RNAscope Probe - Hs-COL4A1 Cat# 461881 and RNAscope Probe - Hs-LAMC2 Cat# 501371. Immunohistochemistry for ActA2 was performed at 1:200 dilution with Abcam Cat# ab5694. The Cy3 filter was used to detect RNA signals and the AF647 filter was used to detect ActA2 protein signal.

### Basement membrane continuity analysis

To evaluate the continuity of BM in the IF images, the BM location was annotated using Qupath (RRID:SCR_018257) (32). The percentage of continuity of the BM for each duct was determined by calculating the ratio of the total number of intact pixels to the total number of pixels in the BM. Each duct was identified by a pathologist-annotated mask that indicates the precise location of the duct on the core. Calculations were performed for the angles of the tangential line to the surface of the mask for each duct in Python. This process aims to offer the algorithm guidance regarding the approximate shape of the duct ensuring a comprehensive scanning of the membrane. With the shape of each duct established, the algorithm evaluates every pixel unit along the membrane to determine if a particular location is deemed intact. This scanning process was conducted within a confidence range of 250-500 pixel units to account for potential variability in annotations.

### WGS DNA-Seq data processing and analyses

DNA sequencing data were preprocessed using BWA-MEM algorithm v0.7.17 (RRID:SCR_010910) (33) for sequence alignment to the reference genome GRCh38/hg38 and GATK v4.1.7.0 (RRID:SCR_001876) (34–36) to mark duplicates and calibrate reads within the Nextflow-based pipeline Sarek v2.6.1 (37,38). The recalibrated reads were further processed and filtered for mappability, GC content using the R/Bioconductor package QDNAseq v1.22.0 (RRID:SCR_003174) (39) with R statistical environment v3.6.0 (RRID:SCR_001905) (40). For QDNAseq, 50-kb bin annotations were obtained from QDNAseq.hg38 (v1.0.0, https://github.com/asntech/QDNAseq.hg38). Only autosomal sequences were retained after filtering based on low-depth mappability and GC correction. The R package ACE v1.4.0 (41) was used to estimate and identify the max cellularity for different ploidies (2, 3, 4). Copy number aberrations were called using CGHcall v2.48.0 (RRID:SCR_001578) (42).

### Bulk RNA-Seq data processing

The RNA sequencing libraries were pre-processed using 3SEQtools (31). Read alignment was conducted using STAR v2.7.3a (RRID:SCR_004463) (43). Gene level reads were obtained using featurecounts function from Rsubread package v4.0.0(RRID:SCR_016945) (44). The reference sequence and genomic annotation files were obtained from GENCODE (RRID:SCR_014966) (45).

### scRNA-seq data processing and analyses

#### Preprocessing

To process the raw scRNA-seq data into a cell by gene matrix with read counts, 10X Genomics Cell Ranger v6.0.1 (RRID:SCR_023221) (46) was used. Briefly, the mkfastq module used bcl2fastq to demultiplex the reads, then the count module aligned and mapped reads to the human reference genome (reference bundle version "refdata-gex-GRCh38-2020-A") and then estimated read counts for each cell across 36,601 annotated genes. The resulting cell-by-gene count matrices were imported into the R v4.2.2 (RRID:SCR_001905) (47) using the Seurat package v4.3.0.1 (RRID:SCR_016341) (48) to perform quality control and downstream analyses.

#### QC

Data quality was assessed in the following ways. Cell quality was assessed by examining the distribution of reads and genes detected per cell and the proportion of reads mapping to mitochondrial genes. A gene was considered to be “detected” if at least one read was mapped to it. Rare gene features were removed from the dataset by excluding genes that were detected in fewer than 10 cells per sample. Mitochondrial genes were identified as features whose corresponding HGNC symbol (RRID:SCR_002827) (49) were prefixed by the string “MT-”. At a minimum, cells were assumed to be low quality and excluded if they had fewer than 200 detected genes or if mitochondrial reads comprised more than 20% of the total reads. In addition, cells with extremely high numbers of reads or genes detected were filtered out based on sample-specific thresholds. The sample-specific thresholds were determined by examining their distributions and removing small shoulders to both sides of the major peak to remove potential dead cells and multiplets.

Eight libraries (3 DCIS and 5 normal) were excluded from downstream analyses due to an insufficient number of high-quality cells (≈1,000 or fewer), and two IBC libraries were excluded due to low sample size. The remaining 20 libraries (7 normal, 13 DCIS) containing 140,322 cells were derived from 14 patients: 5 patients with paired normal and DCIS libraries (10 libraries), 7 patients with DCIS-only libraries (8 libraries, 2 DCIS containing specimens/libraries from the same patient), and 2 patients with normal-only patient libraries (2 libraries).

#### Cell integration

Quality-filtered count matrices from each library were merged into one dataset for downstream analyses and integrated to account for variation among the 8 sequencing batches. Integration was performed using Seurat (RRID:SCR_016341) (48) functions that involve (i) merging sample count matrices by batch then (ii), within each batch counts were log-normalized and the top 2,000 most variable gene features were selected based on variance-stabilizing transformation. Next, (iii) variable gene features across batches, i.e. integration features, were selected from the union of top batch-wise variable features and these were used to reduce the dimensionality of each dataset and identify mutual nearest neighbors, i.e. anchors. Integration returned a single matrix of log-normalized and batch-corrected counts for each cell and integration feature; hereafter, “integrated data”.

#### Clustering

To visualize cell expression profiles across all samples, the integrated data was scaled, reduced in dimension using principal component analysis (PCA) on the top 30 components, and subsequently reduced onto 2-dimensional Uniform Manifold Approximation and Projection (UMAP) coordinates (50) using Seurat (RRID:SCR_016341) (51) functions. Unsupervised clustering of the cells was performed with Seurat (RRID:SCR_016341) (51) functions by constructing a shared nearest-neighbor (SNN) graph (52) based on the top 30 PCA components and then optimizing the standard modularity function based on the Louvain algorithm (53) with the resolution parameter set to 0.5.

#### Cell-type inference

To infer a cell type classification for each cell based on gene expression, the R package scSorter v0.0.2 (54) was used along with a curated set of signature genes (**Supplementary Figure 4A**). The curated signature gene set consisted of genes to profile the 10 major cell types in normal breast tissue, including Basal/Myoep, LumSec, LumHR, B cells, Fibroblasts, Lymphatic, Myeloid, Perivascular, T cells, Vascular (see Supplementary Table 3 from Kumar *et al.* (16)) and 11 epithelial cell substates, including Basal/Myoep, LumSec-basal, LumSec-HLA, LumSec-KIT, LumSec-lac, LumSec-major, LumSec-myo, LumSec-prol, LumHR-active, LumHR-major, LumHR-SCGB (see Supplementary Table 10 from Kumar *et al.* (16)). Specifically, the signature genes for the 3 major epithelial cell states in the former set were replaced by the signature genes for the 11 epithelial cell substates in the latter set. Cell type inference was performed using the top 2,000 most variable genes, excluding rarely expressed genes that are detected in ≤10% of cells. Counts were log-normalized and a scSorter (54) tuning parameter alpha of 0.2 was selected to allow for unknown cells and represent the most common composition of cell types across a range of alpha values.

#### CNV inference

We distinguished the DCIS cells in DCIS libraries (DD) from the non-neoplastic epithelial cells in DCIS libraries (DN) based on their RNA expression-inferred CNV profile. The inferCNV package v1.10.1 (RRID:SCR_021140) (55) was used to infer the copy number states of cells from DCIS samples using scRNA data. To this end, a panel of 51,151 normal epithelial cells from the 7 normal scRNA-seq libraries was constructed to estimate the diploid state as a reference. A cutoff of 0.1 for the minimum average read counts per gene among reference cells, default settings for Hidden Markov Model (HMM) and de-noising filters were used. In addition, the analysis mode of “subclusters” was used to predict CNV at the levels of subpopulations.

WGS CNV profiles were available for eight of the DCIS samples in the final analysis data set. The CNV profiles were initially assessed through a manual review process based on investigators’ knowledge. During this process, specific DNA-inferred CNV regions were identified to serve as decision rules for classifying each individual cell as either a neoplastic or non-neoplastic cell. For a given cell within a scRNA-seq sample, if the number of RNA-inferred CNV regions aligning with the decision rule exceeded a sample-specific threshold, the cell was designated as a neoplastic cell. In cases where scRNA-seq samples lacked a co-located DNA-inferred CNV profile, the set of decision rule regions was derived from the CNV profile of the scRNA-seq sample itself. In total, 17,044 DD and 14,309 DN cells were identified in 13 DCIS libraries and 14,555 NN cells were identified in 7 normal libraries. The decision rules and CNV profiles for the DCIS samples are provided in **Supplementary Table 2 and Supplementary Figure 2C**.

#### DCIS copy-number state phylogenies

To investigate tumor heterogeneity, we examined cell subpopulations within each DCIS sample with different CNV profiles as identified by Leiden clustering (56) implemented within the inferCNV (RRID:SCR_021140) (55) “subcluster” mode. The cell populations, hereafter “CNV subclones" were characterized by neoplastic status and cell state composition. To show the relationship between CNV subclones and infer alteration histories within a DCIS sample, a phylogenetic tree for each sample was constructed by calculating the Euclidean distance between subclones based on gene-level predicted CNV states, building a neighbor-joining tree, and then rooting the tree using an outgroup with no copy-number alterations. The R packages ape v5.7.1 (RRID:SCR_017343) (57) and those within the treedataverse v0.0.1 (58) were used for tree-building and visualization. For each patient, DE analysis was performed to compare each DD subclone against all the other DD subclones using the FindConservedMarkers function from Seurat package v4.3.0.1 (RRID:SCR_016341) (48) in R. For Gene Set Enrichment Analysis (GSEA), all genes were pre-ranked by the log fold-change of the average expression from the Seurat results prior to the enrichment analysis of Hallmark Pathways using the fgsea package v1.24.0 (RRID:SCR_020938) (59). The collection of pathways was acquired from Molecular Signatures Database (MSigDB, RRID:SCR_016863) (60,61) via the msigdbr package v7.5.1 (RRID:SCR_022870) (62). All *p*-values were adjusted to control for multiple testing using the Benjamini-Hochberg procedure (63).

#### Pseudotime trajectory inference

The monocle package v2.26.0 (RRID:SCR_016339) (64–66) was used to infer the pseudotime trajectory of epithelial cell expression. The read count matrix of the epithelial cells was filtered to remove rarely expressed genes (i.e., excluded genes found in equal or less than 100 cells, or showing an expression value less than 0.1), and underwent dimension reduction to 2-dimensions using Discriminative Dimensionality Reduction with Trees (DDRTree) (67) and then cells were ordered in the reduced space using the signature genes used for cell-type inference. The root point of the trajectory was assigned to the Monocle-inferred state that contained the most Basal cells.

#### Pseudo-bulk analysis

To identify differentially expressed genes raw cell-level counts were summed from each patient, cell state, cell substate, and subclone and then DE and GSEA analyses were performed for various comparisons. DE analysis was performed using the DESeq2 package v1.38.3 (RRID:SCR_015687) (68) in R. For GSEA, all genes were pre-ranked by the Wald statistics from the DESeq2 results prior to the enrichment analysis of Hallmark Pathways using the fgsea package v1.24.0 (RRID:SCR_020938) (59). All *p*-values were adjusted to control for multiple testing using the Benjamini-Hochberg procedure (63).

#### Cell type abundance imputation

The CIBERSORTx tool (RRID:SCR_016955) (69) was used to infer the cell type composition within bulk RNA-seq data sets (Sample sets 1-3), using a published single cell data set (16). For each bulk data set, its raw gene count matrix was used as the input, while the external scRNA-seq count matrix with original cell assignments was used to construct the cell type signature matrix. The matrix was randomly down sampled to include a maximum of 3,000 cells per cell type, for the purpose of computing efficiency. The fraction of cells with identical identities showing evidence of gene expression was set to 0. To ensure robust analysis, 100 permutations were performed for *p*-value calculation. S-mode batch correction, designed specifically for scRNA-seq-derived signature matrices, was applied to correct for cross-platform batch effects. The raw imputed cell fractions were then used to calculate the fraction of each epithelial cell type relative to the total fraction of all epithelial cell types.

### Statistical analyses

To compare imputed cell type abundance across sample types in bulk datasets, and to compare the cell type fractions across sample types in scRNA-seq data, the Wilcoxon rank-sum test was used. In the case where bulk data had a patient-matched design (Sample set 1), the Wilcoxon signed-rank test was used instead. To account for multiple testing arising from various cell types, the resulting *p*-values were corrected using the Benjamini-Hochberg method (63). Fisher’s exact test was used to compare the overall fractions of cells of each contrast group, within each cell type. Pearson’s correlation test was used to evaluate the correlation between the cell type abundance estimated from the scRNA-seq data and the corresponding bulk data (Sample set 1) on a per-sample basis.

### Outcome analysis

Patients in sample set 3 were divided into ‘high’ and ‘low’ cell fraction groups based on the median of each respective cell substate. Associations with time to IBC recurrence were quantified using Cox Proportional Hazard model (70). Kaplan-Meier plots as implemented in the R packages survival v3.3-1 (RRID:SCR_021137) (70) and survminer v0.4.9 (RRID:SCR_021094) (71) were used to visualize outcome differences.

### *In vitro* invasion assay

#### Cell culture

MCF10A human mammary epithelial cells (ATCC, RRID:CVCL_0598) were cultured in Dulbecco’s modified Eagle’s medium/Nutrient Mixture F-12 (DMEM/F12) medium (Thermo Fisher) supplemented with 5% horse serum (Thermo Fisher), 20 ng/ml EGF (Peprotech, Inc.), 0.5 µg/ml hydrocortisone (Sigma), 100 ng/ml cholera toxin (Sigma), 10 µg/ml insulin (Sigma), and 1% penicillin/streptomycin (Thermo Fisher). Cells were passaged every 3-4 days with 0.05% trypsin/EDTA and cultured in a standard humidified incubator at 37°C and 5% CO_2_.

#### Acini formation

Mammary acini using MCF10A mammary epithelial cells were generated as previously described (23). Briefly, single cell suspension of MCF10A cells were seeded onto reconstituted basement membrane (rBM) in 2 mL of growth media supplemented with 2% rBM (72). After 4 days, the media was replenished with rBM supplemented media. On Day 5, acini were extracted by treatment with 50mM EDTA in PBS followed by cell scrapping. After a 20-minute incubation on ice, the acini-containing EDTA mixture was spun down for 5 minutes at 500*g* at 4°C, resuspended in the growth media and centrifuged at 500*g* for another 5 min. After centrifugation, the supernatant was aspirated, and acini were resuspended in DMEM/F12 media.

#### Alginate preparation and acini encapsulation in hydrogels

High molecular weight sodium alginate was synthesized and used to develop interpenetrating hydrogels with specific mechanical stiffness (2.4 kPa) for acini culture as described previously (23).

#### Immunofluorescence of acini

Acini were fixed in 4% paraformaldehyde in serum-free DMEM/F12 at room temperature. After fixation, acini were washed twice in PBS for 15 min and stained with antibodies. The following antibodies were used for detection: Alexa Fluor 488 conjugated anti-laminin-5 antibody, clone D4B5 (Millipore Sigma, MAB19562X,1:200 dilution, RRID:AB_570380), anti-collagen-IV mouse antibody (Sigma, #SAB4500369, 1:200, RRID:AB_10743858) and Alexa Fluor 690 goat anti-mouse (#A21240, 1:1000, RRID:AB_2535809) and nuclear stain Hoechst 33342 (#H3570, 1:1000).

#### Collagenase-IV treatment

Collagenase-IV powder (Sigma #17104019) was reconstituted in Hank’s Balanced Salt Solution to a stock concentration of 25 U/mL and acini were treated with a 1:1000 dilution for 1 hour. Acini were encapsulated in hydrogel immediately after the treatment.

#### Measurement of basement membrane thickness

Confocal images of laminin and collagen-IV were overlaid and distance measured from the inner surface of laminin to the outer surface of collagen-IV at three random locations in the equatorial cross-section and these measurements were averaged to obtain data for each acini in the experiment.

### Confocal microscopy

Microscope imaging was performed with a laser-scanning Leica SP8 confocal microscope (RRID:SCR_018169) or a Nikon Ti2-E inverted microscope (RRID:SCR_023803), both fitted with a temperature and incubator control suitable for live imaging (37°C, 5% CO2). For capturing fluorescent images, the Leica microscope used a 25x NA 0.91 water objective. For capturing phase contrast images of acini, the Nikon microscope used a 10x NA 0.45 dry objective.

## Data availability

The scRNA-seq data, WGS data, bulk RNA sequencing datasets and imaging data have been deposited via the Human Tumor Atlas Network (HTAN) dbGaP Study Accession: phs002371.v1.p1 (https://www.ncbi.nlm.nih.gov/projects/gap/cgi-bin/study.cgi?study_id=phs002371.v1.p1).

## Code availability

The scripts to reproduce the analyses presented in this paper are available publicly through a gitlab repository (https://gitlab.oit.duke.edu/dcibioinformatics/pubs/pca-dcis-scrna-seq).

## Acknowledgements

Research reported in this publication was supported by National Cancer Institute of the National Institutes of Health under award number U2C CA-17-035 Pre-Cancer Atlas (PCA) Research Centers (E.S.H., M.R.L, J.R.M, K.O., S.H.S., R.B.W., X.Q.) and CA014236 (M.R.L, K.O., X.Q.).

The content is solely the responsibility of the authors and does not necessarily represent the official views of the National Institutes of Health.

## Additional information

Supplementary Information is available for this paper: Supplementary File 1.

Correspondence and requests for materials should be addressed to Dr. Robert B. West.

---

**Supplementary Figure 1. QC plots.**

QC plots for all scRNA-seq samples before and after filtering. A) Distribution of the number of molecules detected per cell. B) Number of genes detected per cell. C) Percentage pf mitochondrial genes per cell. D) Relationship between the number of molecules and the number of genes per cell, colored by percentage of mitochondrial genes. E) Total number of cells.

**Supplementary Figure 2.**

**A)** Distribution of 11 mammary epithelial cell substates in each sample, with text in each cell indicating the raw cell count at the top and the cell count normalized by library size in parentheses. **B)** Histology of specimens used for scRNAseq. Scanned hematoxylin and eosin stained section from the bisected facing half of the specimen dispersed for single cell analysis. Sections were scanned at 20X magnification and these screenshots from the scans show each entire specimen. **C)** Heatmaps of single cells with copy number status at each genomic locus, inferred from a low-pass WGS (upper panel) and inferred from single-cell DCIS (middle panel) libraries. A panel of 7 normal libraries were used as the reference (bottom panel). Each page shows an individual DCIS specimen. **D)** GSEA of all 50 Hallmark pathways between pseudo-bulk samples from neoplastic and non-neoplastic epithelial cells from breast tissue samples. The numbers of significant genes are indicated on top of the heatmap of the different comparisons. The color of the dots indicates the normalized enrichment score, and the size indicates significance with the negative log_10_ adjusted *p*-value.

**Supplementary Figure 3. CNV phylogenetic trees.**

**A)** Heterogeneity of CNV profiles in DCIS samples from each patient. For each of 12 patients, we show the subclone CNV phylogenetic trees with neoplastic status classifications as shown by tree tip and subclone label colors. The number of cells, cell state/substate compositions, and inferred CNV regions by subclone are shown in the bar charts and tile plots. **B)** GSEA results for a given DD subclone as compared to all the other DDs in the same patient DCIS tissue. Of the 12 patients, 9 had more than one DD subclone which was necessary to perform this analysis and generate the results shown here. **C)** and **D)** Position of DCIS genetic subclones in **C)** Pseudotime trajectory space and **D)** UMAP space. Subclones are colored separately for each case and reference to the phylogenies indicate the identity of the subclones. The pseudotime trajectory (**Figure 3I**) and UMAP (**Figure 3A**) were constructed from all scRNA-seq epithelial cells. **E)** Inferred CNV state at *ERBB2* in DCIS subclones from each patient. Subclone neoplastic status is shown by point color and the patient’s ER status is indicated in the panel headers.

**Supplementary Figure 4. Mammary cell state and substate analysis.**

**A)** Genes used to define cell types and mammary epithelial cell states and substates. **B)** GSEA of all 50 Hallmark pathways between pseudo-bulk samples from DN/NN and DD of three major cell states in ER-DCIS and ER+ DCIS, analyzed separately. **C)** GSEA analysis as in **B)** but for the 11 mammary substates. The color of the dots indicates the normalized enrichment score, and the size indicates significance with the negative log_10_ adjusted *p*-value.

**Supplementary Figure 5. Bulk sample cell fraction analysis.**

**A)** Correlation between relative cell fractions from the 13 pseudo-bulk samples (5 DD ER-, 6 DD ER+, and 2 DNNN) derived from the 12 scRNA-seq specimens (10 DCIS, 2 Normal) and imputed relative cell fractions from 13 patient-matched FFPE bulk RNA-seq samples (5 ER-DCIS, 6 ER+ DCIS, and 2 normal), across 3 major epithelial cell states (Sample set 1). **B)** Analysis of imputed cell fractions based on 11 epithelial substates for 163 DCIS that did not progress or recur and 69 DCIS that progressed to IBC from Sample set 3.

**Supplementary Figure 6. Expression of basement membrane genes.**

**A)** Relative expression of BM genes across 11 epithelial cell substates and fibroblasts within each pseudo-bulk sample from scRNA-seq data (Sample set 1). Displayed genes were most significantly up-regulated (adj. *p* < 0.01) in each specific cell type compared to all other cell types. **B)** Immunofluorescent staining of COL4A1 (red) in normal breast and **C)** LAMC2 (green) in normal breast with blue DAPI nuclear stain (Sample set 4).

