## Supplementary Figures for "Single Cell Expression Analysis of Ductal Carcinoma in Situ Identifies Complex Genotypic-Phenotypic Relationships Altering Epithelial Composition"

### Supplementary Figure 1

## A

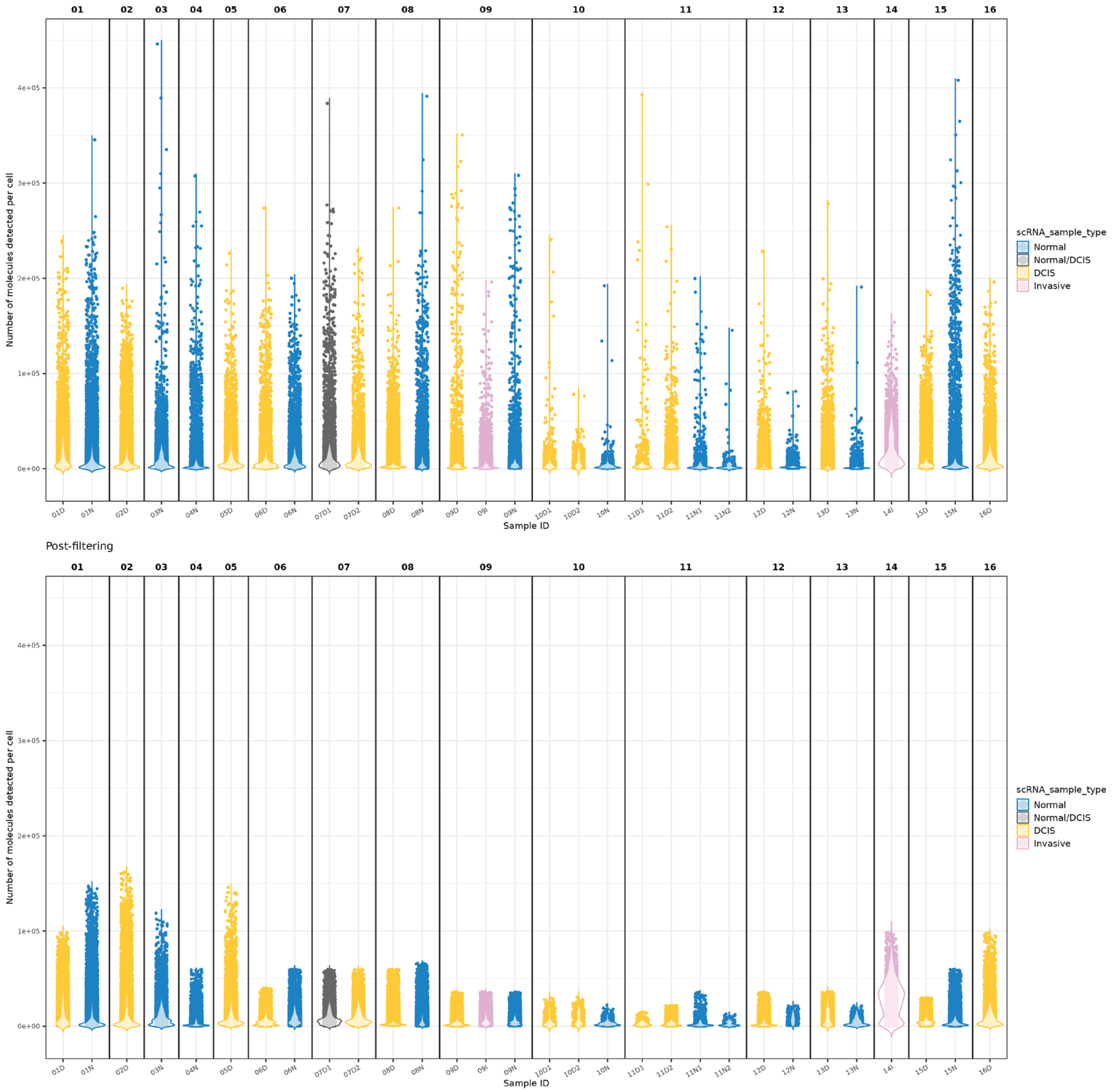

### Supplementary Figure 1 (Cont.)

## B

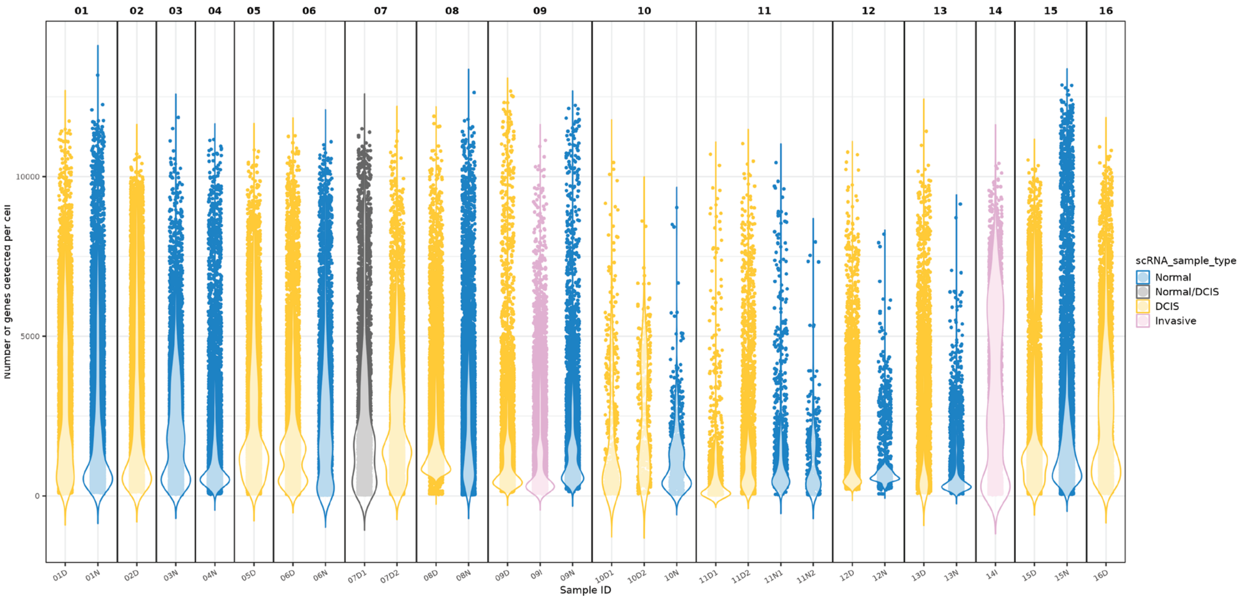


Post-filtering

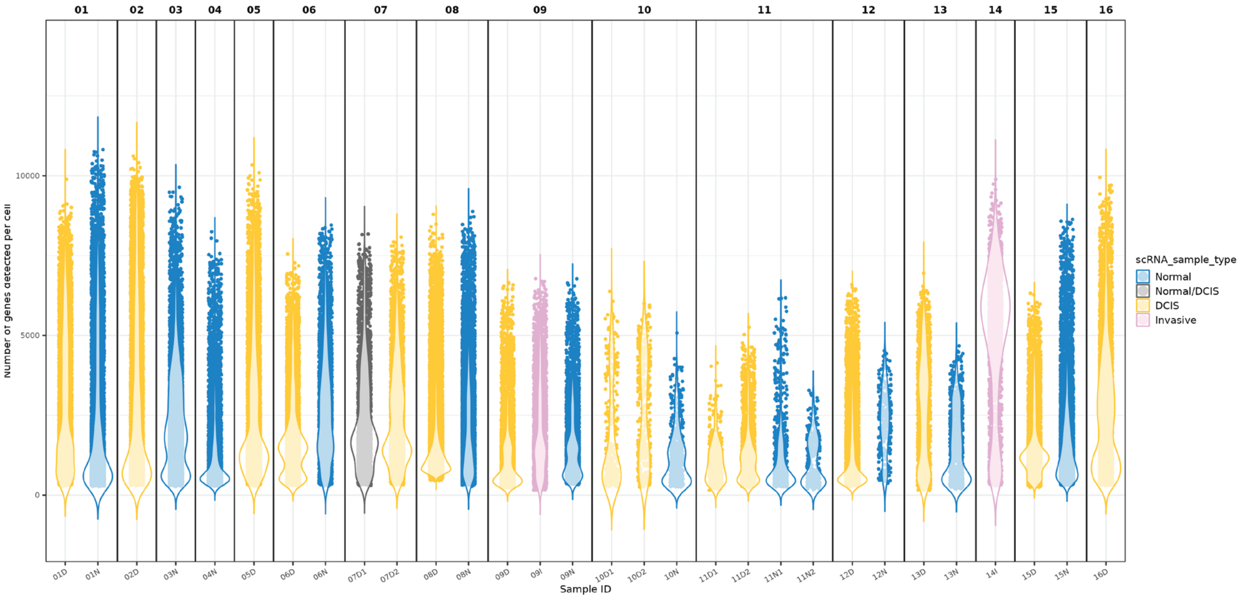

### Supplementary Figure 1 (Cont.)

C

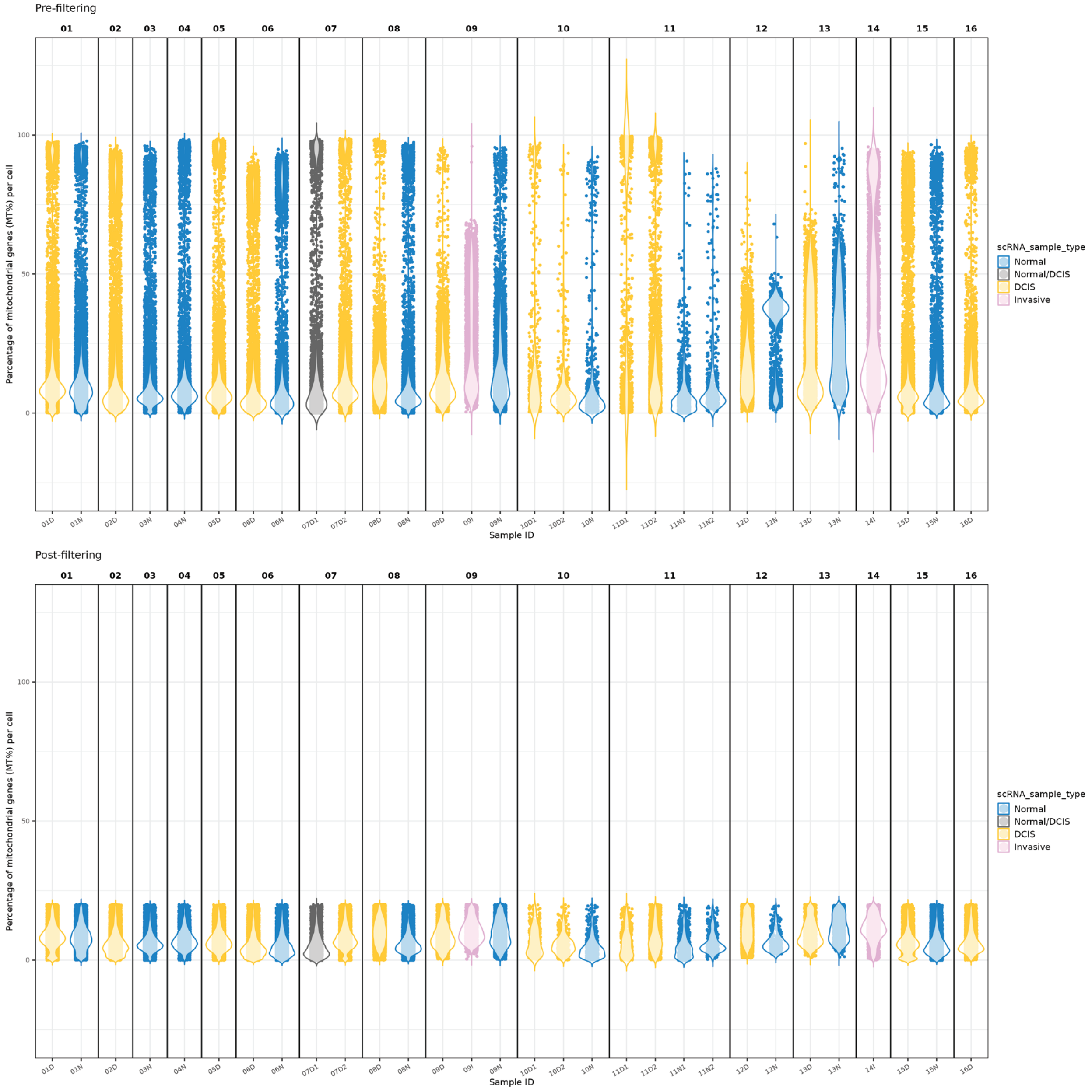

### Supplementary Figure 1 (Cont.)
## D

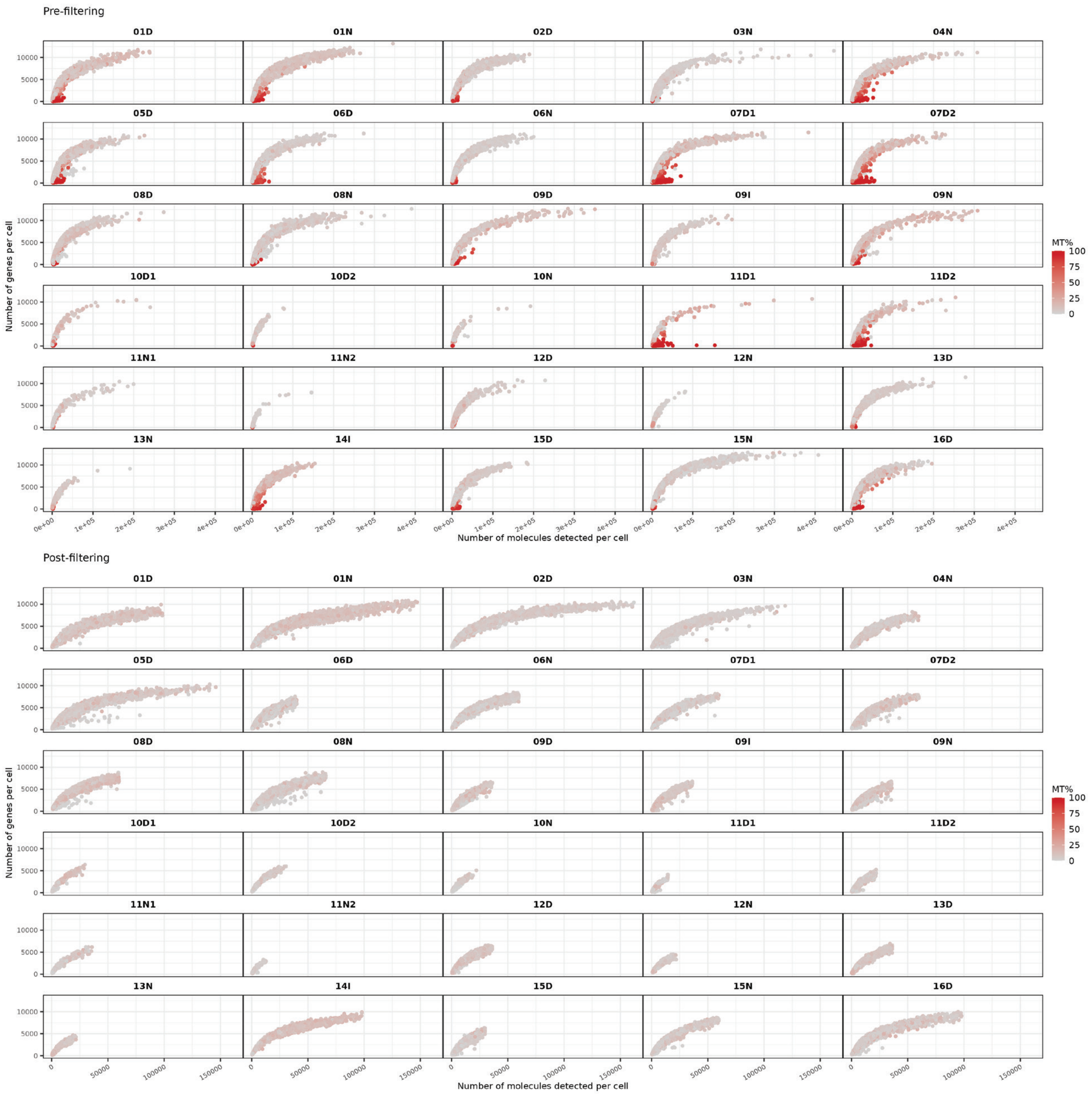

#### Supplementary Figure 1 (Cont.)

## E

| | Sample ID | Pre-filtering | Post-filtering | Cells Kept% |
|---|---|---|---|---|
| 1 | 01D | 9964 | 7886 | 79.14492 |
| 2 | 01N | 10611 | 8681 | 81.81133 |
| 3 | 02D | 12903 | 10310 | 79.90390 |
| 4 | 03N | 10835 | 9359 | 86.37748 |
| 5 | 04N | 7326 | 6066 | 82.80098 |
| 6 | 05D | 9900 | 8626 | 87.13131 |
| 7 | 06D | 9478 | 7396 | 78.03334 |
| 8 | 06N | 6031 | 4455 | 73.86835 |
| 9 | 07D1 | 4750 | 3235 | 68.10526 |
| 10 | 07D2 | 7773 | 6197 | 79.72469 |
| 11 | 08D | 19361 | 17841 | 92.14917 |
| 12 | 08N | 9428 | 8082 | 85.72338 |
| 13 | 09D | 6162 | 4983 | 80.86660 |
| 14 | 09I | 6972 | 3481 | 49.92828 |
| 15 | 09N | 6950 | 5026 | 72.31655 |
| 16 | 10D1 | 538 | 371 | 68.95911 |
| 17 | 10D2 | 496 | 428 | 86.29032 |
| 18 | 10N | 762 | 635 | 83.33333 |
| 19 | 11D1 | 1323 | 438 | 33.10658 |
| 20 | 11D2 | 3576 | 2194 | 61.35347 |
| 21 | 11N1 | 1114 | 955 | 85.72711 |
| 22 | 11N2 | 470 | 367 | 78.08511 |
| 23 | 12D | 9675 | 7161 | 74.01550 |
| 24 | 12N | 1971 | 297 | 15.06849 |
| 25 | 13D | 4672 | 2009 | 43.00086 |
| 26 | 13N | 2623 | 1046 | 39.87800 |
| 27 | 14I | 5363 | 2343 | 43.68823 |
| 28 | 15D | 7482 | 5030 | 67.22801 |
| 29 | 15N | 10981 | 9482 | 86.34915 |
| 30 | 16D | 7358 | 6303 | 85.66186 |

**Supplementary Figure 1. QC plots.**
QC plots for all scRNA-seq samples before and after filtering. **A)** Distribution of the number of molecules detected per cell. **B)** Number of genes detected per cell. **C)** Percentage pf mitochondrial genes per cell. **D)** Relationship between the number of molecules and the number of genes per cell, colored by percentage of mitochondrial genes. **E)** Total number of cells.

### Supplementary Figure 2

## A

Sample Type: Normal

| Cell type | 01N | 03N | 04N | 06N | 08N | 09N | 15N |
|---|---|---|---|---|---|---|---|
| Basal/Myoep | 51 (0.006) | 27 (0.003) | 93 (0.015) | 23 (0.005) | 133 (0.016) | 190 (0.038) | 265 (0.028) |
| LumSec−basal | 342 (0.039) | 150 (0.016) | 354 (0.058) | 666 (0.149) | 967 (0.120) | 9 (0.002) | 252 (0.027) |
| LumSec−HLA | 36 (0.004) | 574 (0.061) | 445 (0.073) | 0 (0.000) | 0 (0.000) | 0 (0.000) | 0 (0.000) |
| LumSec−KIT | 149 (0.017) | 90 (0.010) | 124 (0.020) | 173 (0.039) | 61 (0.008) | 35 (0.007) | 204 (0.022) |
| LumSec−lac | 0 (0.000) | 5 (0.001) | 0 (0.000) | 1 (0.000) | 4 (0.000) | 218 (0.043) | 5 (0.001) |
| LumSec−major | 18 (0.002) | 1 (0.000) | 6 (0.001) | 15 (0.003) | 113 (0.014) | 1,682 (0.335) | 135 (0.014) |
| LumSec−myo | 1 (0.000) | 0 (0.000) | 0 (0.000) | 0 (0.000) | 6 (0.001) | 0 (0.000) | 18 (0.002) |
| LumSec−prol | 62 (0.007) | 61 (0.007) | 96 (0.016) | 19 (0.004) | 1 (0.000) | 12 (0.002) | 13 (0.001) |
| LumHR−active | 18 (0.002) | 85 (0.009) | 58 (0.010) | 0 (0.000) | 62 (0.008) | 42 (0.008) | 431 (0.045) |
| LumHR−major | 237 (0.027) | 83 (0.009) | 244 (0.040) | 640 (0.144) | 408 (0.050) | 13 (0.003) | 152 (0.016) |
| LumHR−SCGB | 2,898 (0.334) | 10 (0.001) | 12 (0.002) | 412 (0.092) | 364 (0.045) | 126 (0.025) | 355 (0.037) |
| B cells | 26 (0.003) | 68 (0.007) | 124 (0.020) | 9 (0.002) | 323 (0.040) | 23 (0.005) | 43 (0.005) |
| Fibroblasts | 638 (0.073) | 2,828 (0.302) | 1,308 (0.216) | 499 (0.112) | 1,805 (0.223) | 969 (0.193) | 2,438 (0.257) |
| Lymphatic | 57 (0.007) | 35 (0.004) | 157 (0.026) | 163 (0.037) | 792 (0.098) | 24 (0.005) | 204 (0.022) |
| Myeloid | 1,978 (0.228) | 1,492 (0.159) | 855 (0.141) | 551 (0.124) | 600 (0.074) | 125 (0.025) | 1,220 (0.129) |
| Perivascular | 164 (0.019) | 322 (0.034) | 209 (0.034) | 41 (0.009) | 197 (0.024) | 72 (0.014) | 309 (0.033) |
| T cells | 1,121 (0.129) | 2,098 (0.224) | 408 (0.067) | 948 (0.213) | 1,287 (0.159) | 938 (0.187) | 1,627 (0.172) |
| Vascular | 400 (0.046) | 1,047 (0.112) | 1,184 (0.195) | 213 (0.048) | 778 (0.096) | 407 (0.081) | 1,531 (0.161) |
| Unknown | 485 (0.056) | 383 (0.041) | 389 (0.064) | 82 (0.018) | 181 (0.022) | 141 (0.028) | 280 (0.030) |

Sample Type: DCIS

| Cell type | 01D | 02D | 05D | 06D | 07D1 | 07D2 | 08D | 09D | 11D2 | 12D | 13D | 15D | 16D |
|---|---|---|---|---|---|---|---|---|---|---|---|---|---|
| Basal/Myoep | 38 (0.005) | 84 (0.008) | 55 (0.006) | 32 (0.004) | 18 (0.006) | 11 (0.002) | 110 (0.006) | 282 (0.057) | 86 (0.039) | 851 (0.119) | 29 (0.014) | 25 (0.005) | 100 (0.016) |
| LumSec−basal | 160 (0.020) | 467 (0.045) | 454 (0.053) | 500 (0.068) | 116 (0.036) | 55 (0.009) | 126 (0.007) | 13 (0.003) | 488 (0.222) | 929 (0.130) | 23 (0.011) | 154 (0.031) | 500 (0.079) |
| LumSec−HLA | 16 (0.002) | 3,389 (0.329) | 4 (0.000) | 1 (0.000) | 0 (0.000) | 0 (0.000) | 25 (0.001) | 1 (0.000) | 0 (0.000) | 6 (0.001) | 36 (0.018) | 0 (0.000) | 0 (0.000) |
| LumSec−KIT | 74 (0.009) | 225 (0.022) | 117 (0.014) | 485 (0.066) | 34 (0.011) | 28 (0.005) | 66 (0.004) | 55 (0.011) | 5 (0.002) | 468 (0.065) | 23 (0.011) | 129 (0.026) | 239 (0.038) |
| LumSec−lac | 13 (0.002) | 2 (0.000) | 3 (0.000) | 7 (0.001) | 1 (0.000) | 1 (0.000) | 32 (0.002) | 194 (0.039) | 11 (0.005) | 50 (0.007) | 4 (0.002) | 0 (0.000) | 4 (0.001) |
| LumSec−major | 16 (0.002) | 10 (0.001) | 23 (0.003) | 99 (0.013) | 4 (0.001) | 28 (0.005) | 36 (0.002) | 1,749 (0.351) | 8 (0.004) | 485 (0.068) | 30 (0.015) | 14 (0.003) | 186 (0.030) |
| LumSec−myo | 0 (0.000) | 9 (0.001) | 1 (0.000) | 0 (0.000) | 0 (0.000) | 0 (0.000) | 2,196 (0.123) | 0 (0.000) | 0 (0.000) | 0 (0.000) | 2 (0.001) | 0 (0.000) | 1 (0.000) |
| LumSec−prol | 133 (0.017) | 105 (0.010) | 198 (0.023) | 456 (0.062) | 85 (0.026) | 37 (0.006) | 22 (0.001) | 12 (0.002) | 145 (0.066) | 14 (0.002) | 49 (0.024) | 292 (0.058) | 94 (0.015) |
| LumHR−active | 58 (0.007) | 319 (0.031) | 0 (0.000) | 2 (0.000) | 0 (0.000) | 0 (0.000) | 5,287 (0.296) | 59 (0.012) | 0 (0.000) | 2 (0.000) | 79 (0.039) | 0 (0.000) | 72 (0.011) |
| LumHR−major | 159 (0.020) | 228 (0.022) | 243 (0.028) | 308 (0.042) | 47 (0.015) | 59 (0.010) | 187 (0.010) | 39 (0.008) | 10 (0.005) | 215 (0.030) | 331 (0.165) | 171 (0.034) | 70 (0.011) |
| LumHR−SCGB | 1,486 (0.188) | 48 (0.005) | 675 (0.078) | 857 (0.116) | 79 (0.024) | 39 (0.006) | 55 (0.003) | 167 (0.034) | 30 (0.014) | 1,361 (0.190) | 164 (0.082) | 240 (0.048) | 214 (0.034) |
| B cells | 86 (0.011) | 9 (0.001) | 119 (0.014) | 733 (0.099) | 113 (0.035) | 254 (0.041) | 705 (0.040) | 37 (0.007) | 66 (0.030) | 108 (0.015) | 11 (0.005) | 182 (0.036) | 81 (0.013) |
| Fibroblasts | 458 (0.058) | 1,128 (0.109) | 871 (0.101) | 186 (0.025) | 456 (0.141) | 1,668 (0.269) | 3,118 (0.175) | 509 (0.102) | 26 (0.012) | 272 (0.038) | 445 (0.222) | 150 (0.030) | 1,271 (0.202) |
| Lymphatic | 60 (0.008) | 114 (0.011) | 240 (0.028) | 59 (0.008) | 31 (0.010) | 117 (0.019) | 280 (0.016) | 39 (0.008) | 8 (0.004) | 58 (0.008) | 15 (0.007) | 16 (0.003) | 18 (0.003) |
| Myeloid | 2,753 (0.349) | 886 (0.086) | 1,013 (0.117) | 906 (0.122) | 461 (0.143) | 1,054 (0.170) | 788 (0.044) | 153 (0.031) | 215 (0.098) | 711 (0.099) | 126 (0.063) | 975 (0.194) | 635 (0.101) |
| Perivascular | 96 (0.012) | 361 (0.035) | 235 (0.027) | 20 (0.003) | 212 (0.066) | 216 (0.035) | 215 (0.012) | 109 (0.022) | 55 (0.025) | 92 (0.013) | 97 (0.048) | 65 (0.013) | 577 (0.092) |
| T cells | 1,341 (0.170) | 868 (0.084) | 2,802 (0.325) | 2,047 (0.277) | 851 (0.263) | 1,218 (0.197) | 481 (0.027) | 912 (0.183) | 820 (0.374) | 534 (0.075) | 125 (0.062) | 2,027 (0.403) | 902 (0.143) |
| Vascular | 511 (0.065) | 1,479 (0.143) | 1,236 (0.143) | 189 (0.026) | 640 (0.198) | 1,357 (0.219) | 1,611 (0.090) | 498 (0.100) | 119 (0.054) | 226 (0.032) | 282 (0.140) | 294 (0.058) | 1,103 (0.175) |
| Unknown | 428 (0.054) | 579 (0.056) | 337 (0.039) | 509 (0.069) | 87 (0.027) | 55 (0.009) | 2,501 (0.140) | 155 (0.031) | 102 (0.046) | 779 (0.109) | 138 (0.069) | 296 (0.059) | 236 (0.037) |

**Sample Type**
Normal
DCIS

**Patient**
01
02
03
04
05
06
07
08
09
11
12
13
15
16

**% Normalized Cell Count By Row**
100
50
0

## B

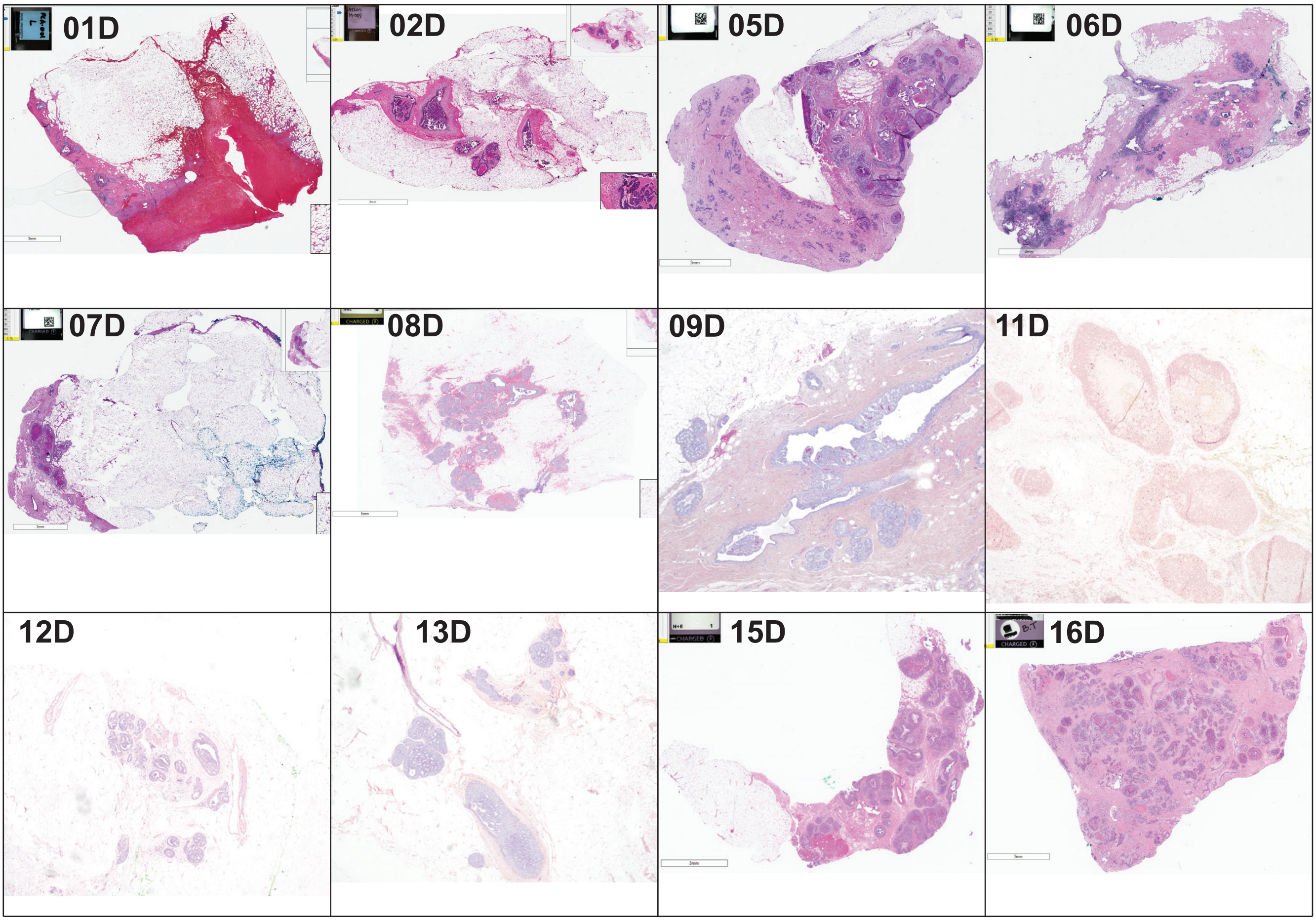

#### Supplementary Figure 2 (Cont.)

C

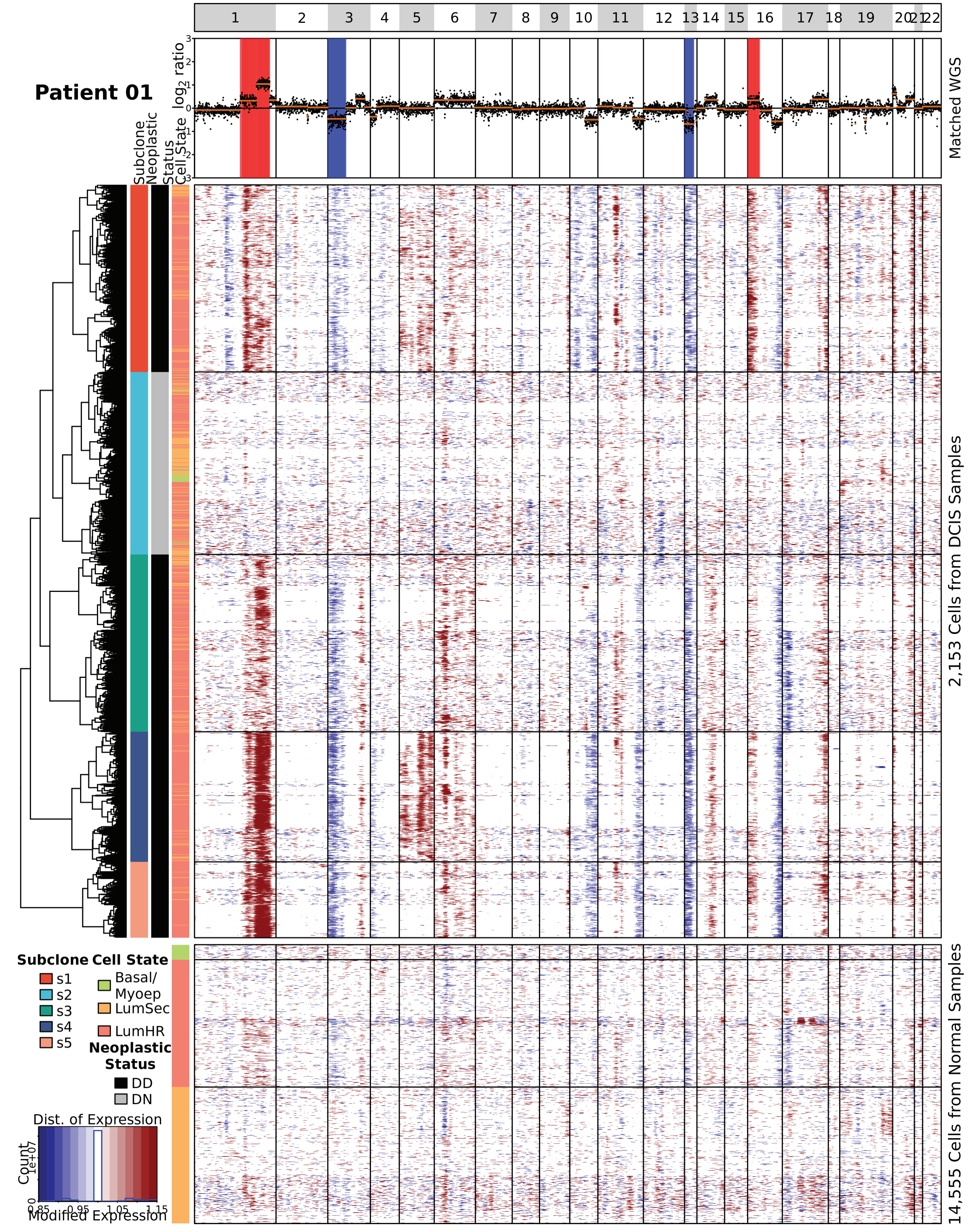

#### Supplementary Figure 2 (Cont.)

**C**

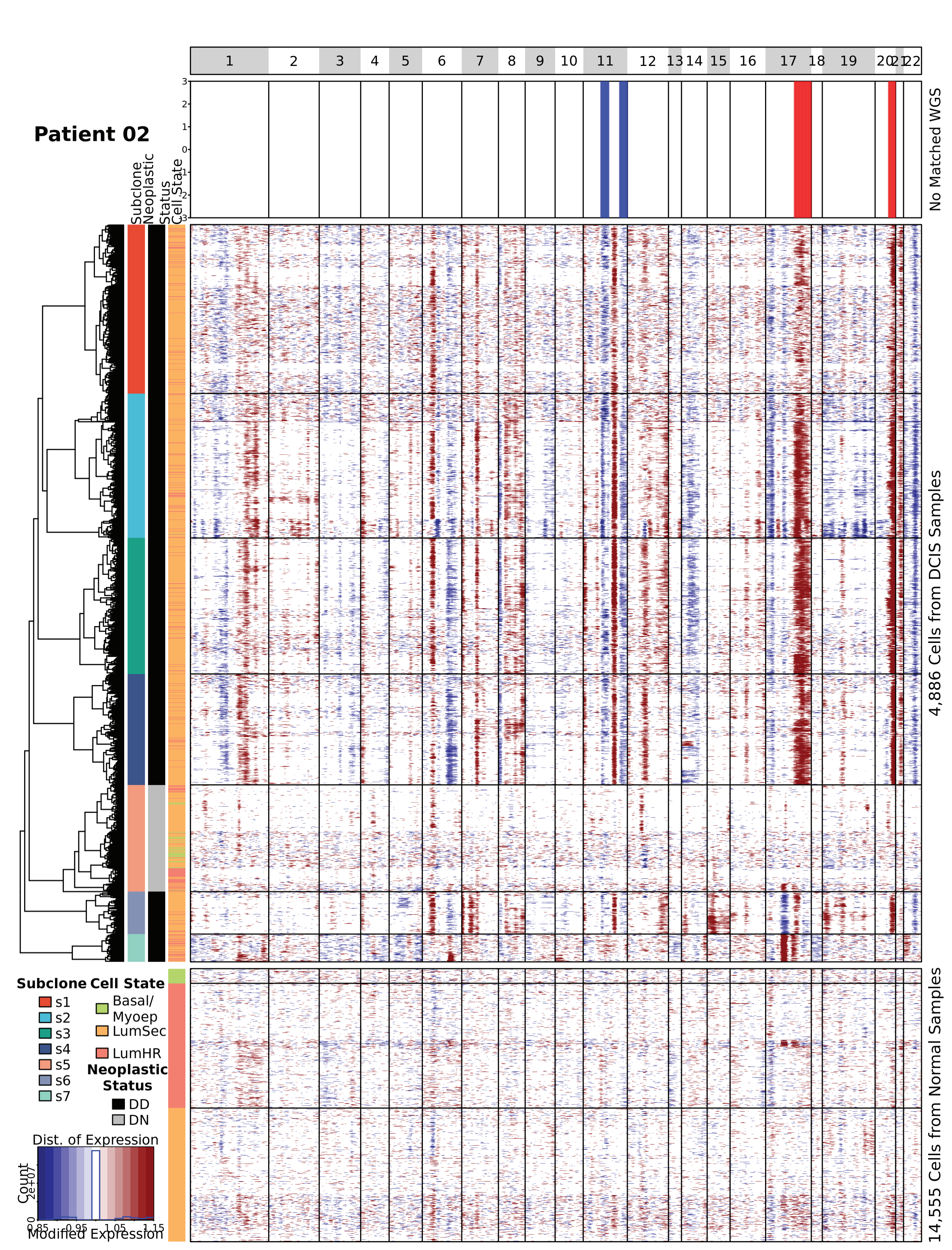

#### Supplementary Figure 2 (Cont.)

**C**

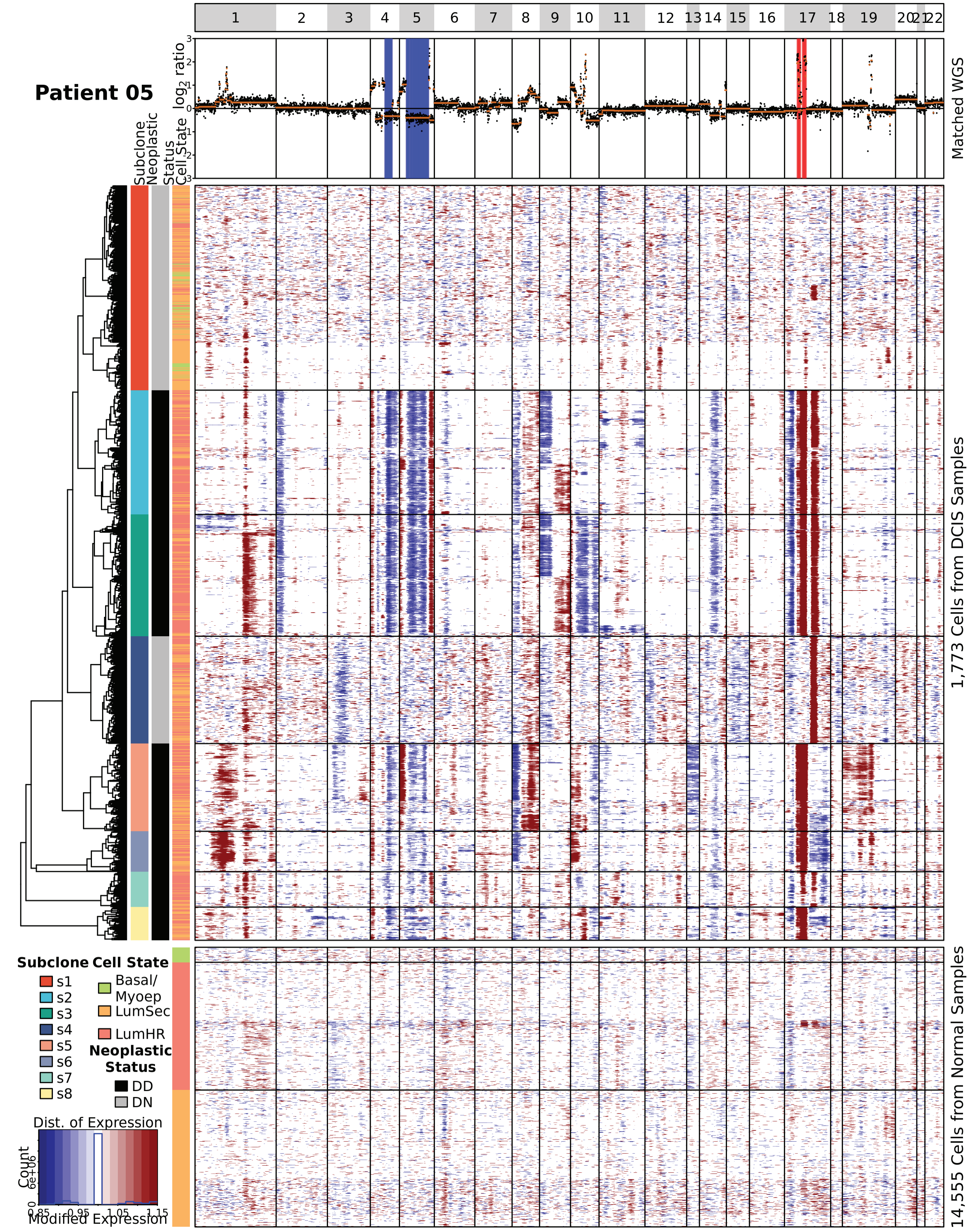

### Supplementary Figure 2 (Cont.)

C

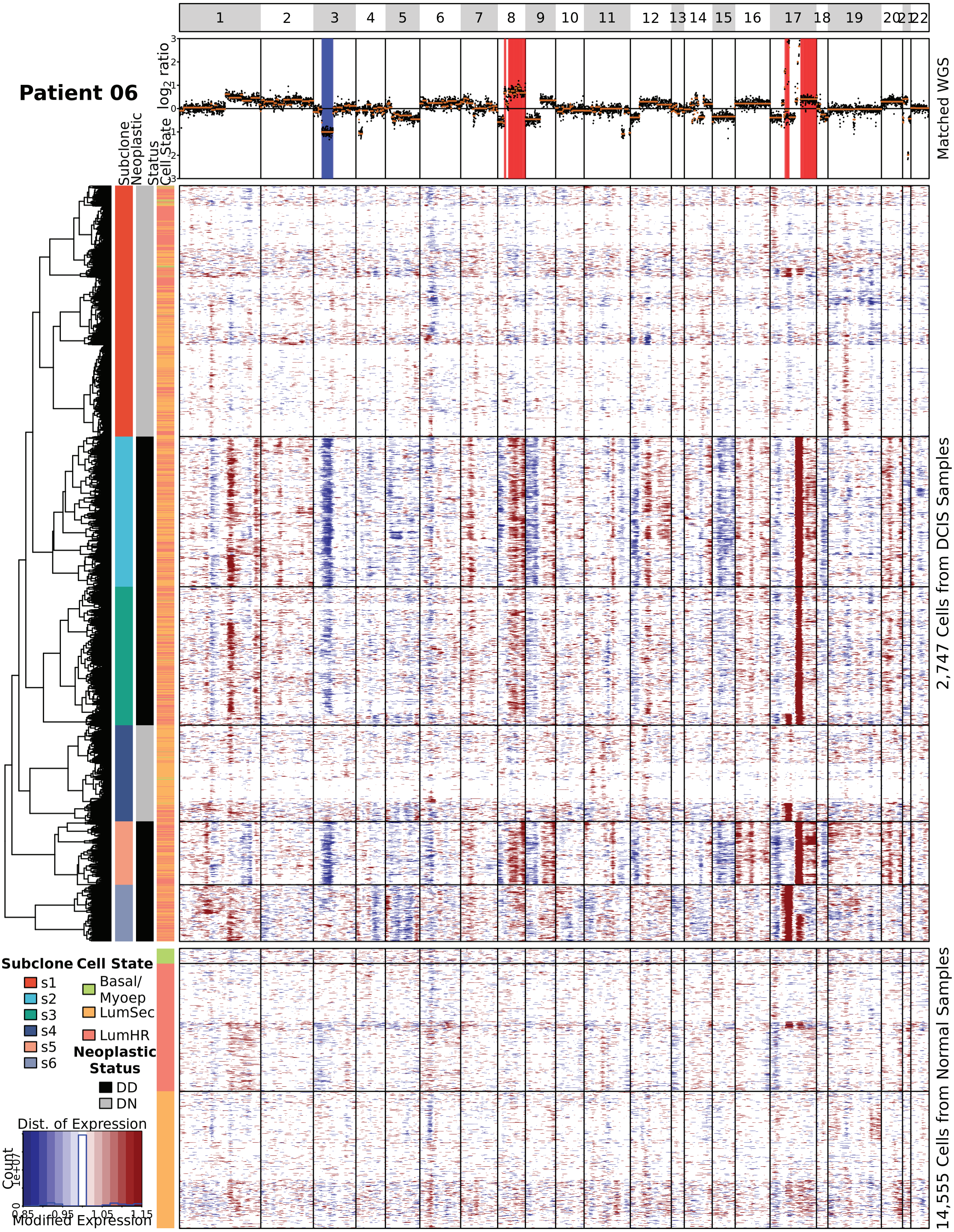

#### Supplementary Figure 2 (Cont.)

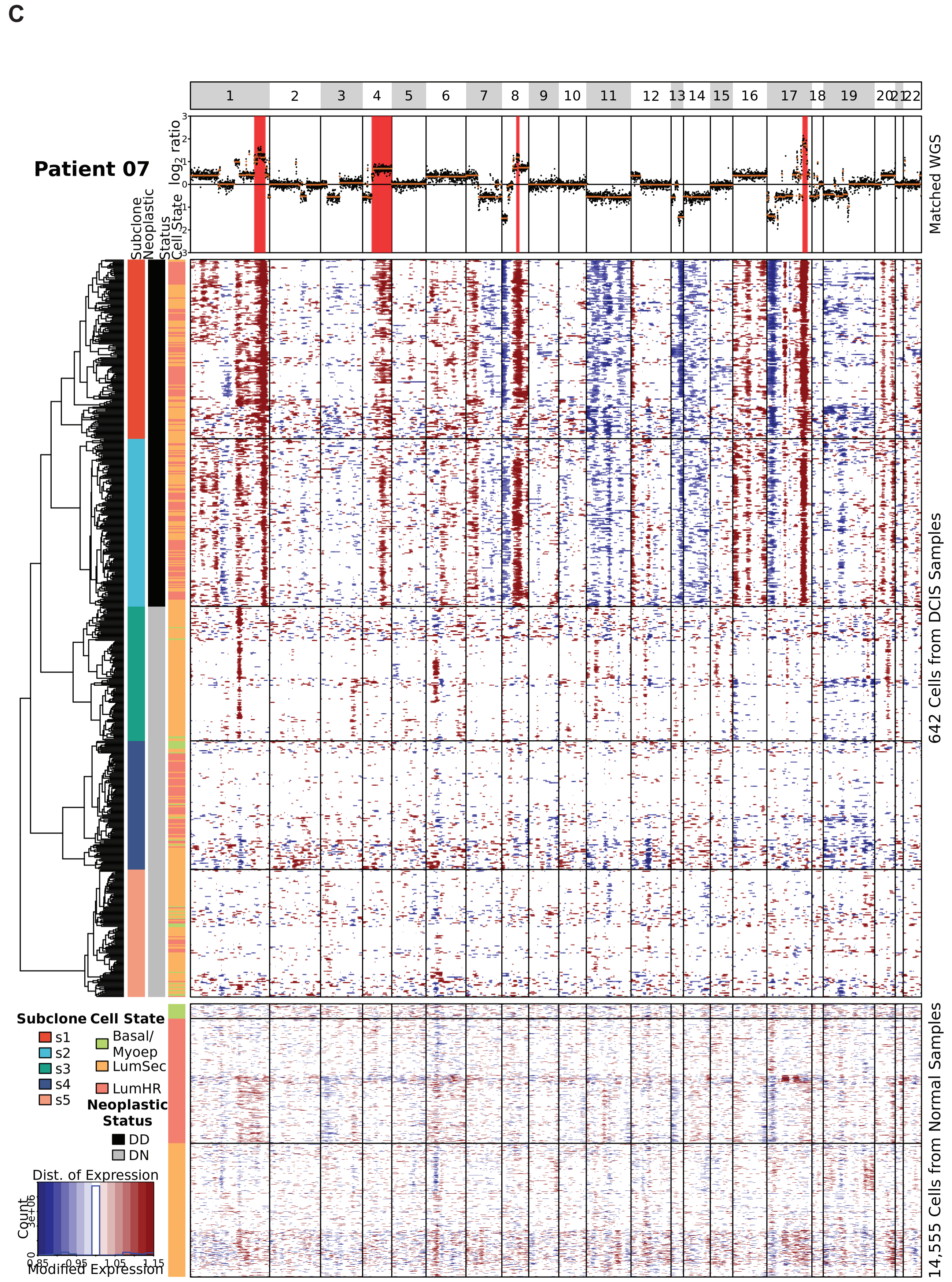

Supplementary Figure 2 (Cont.)

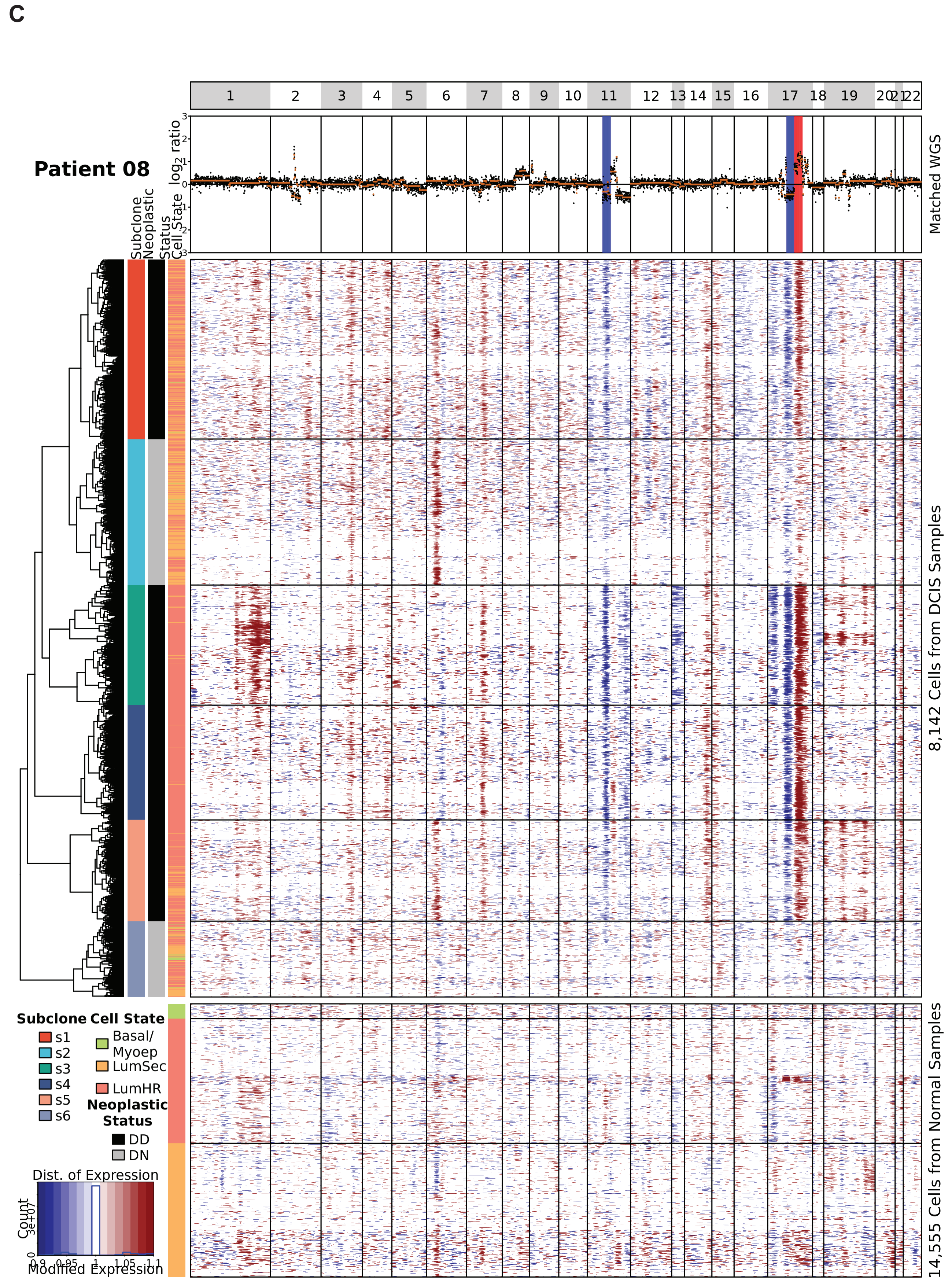

#### Supplementary Figure 2 (Cont.)

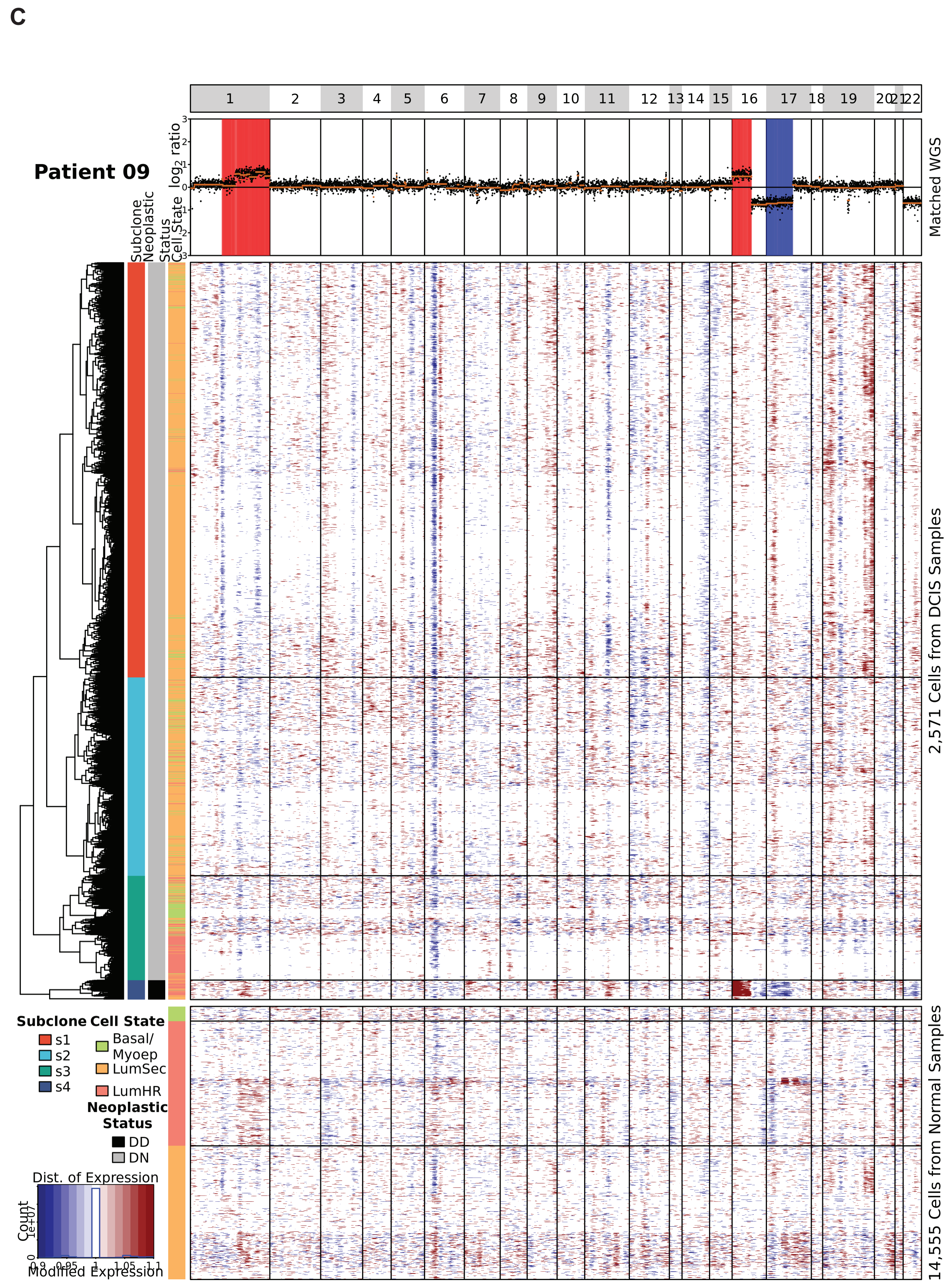

Supplementary Figure 2 (Cont.)

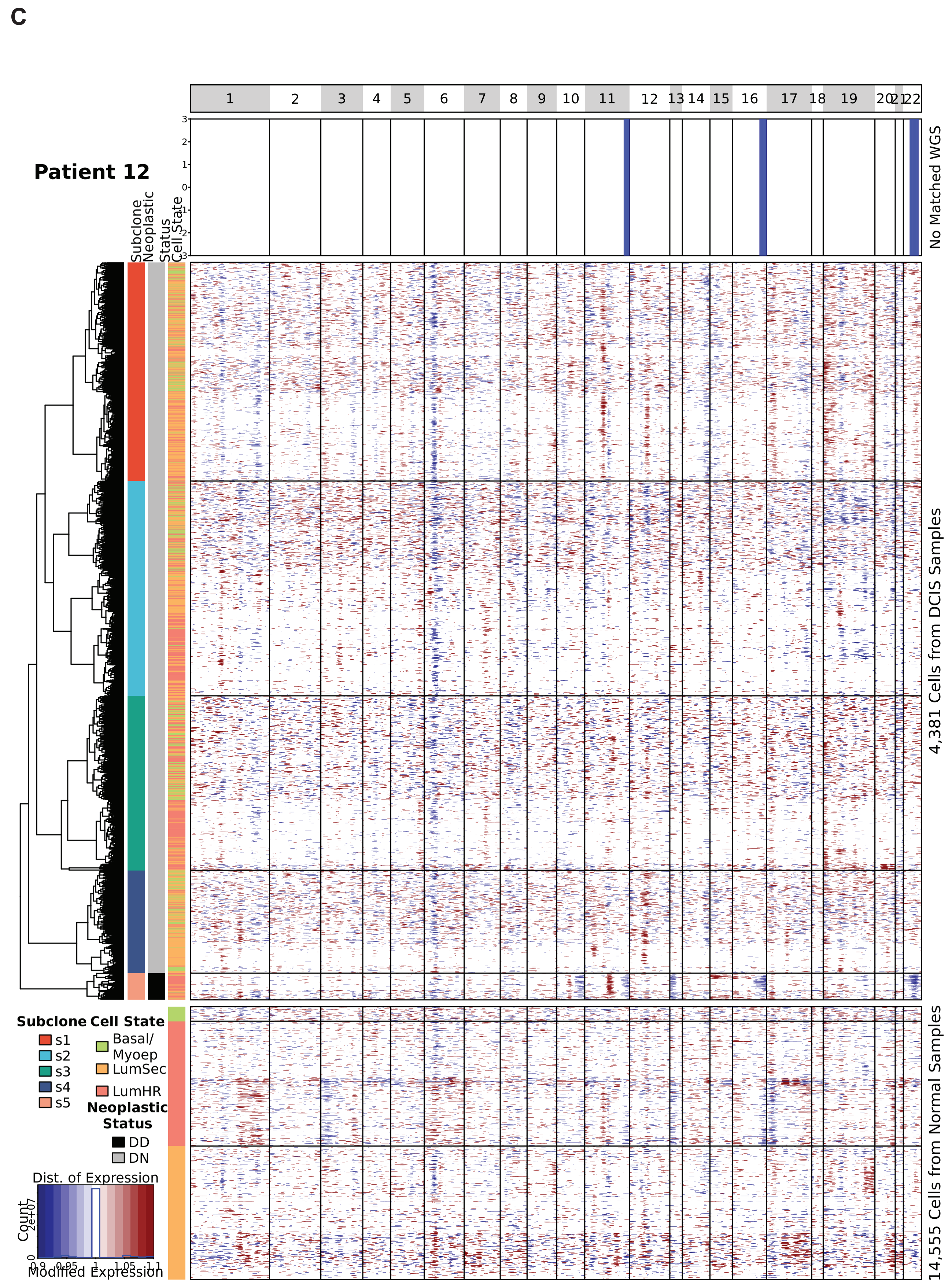

#### Supplementary Figure 2 (Cont.)

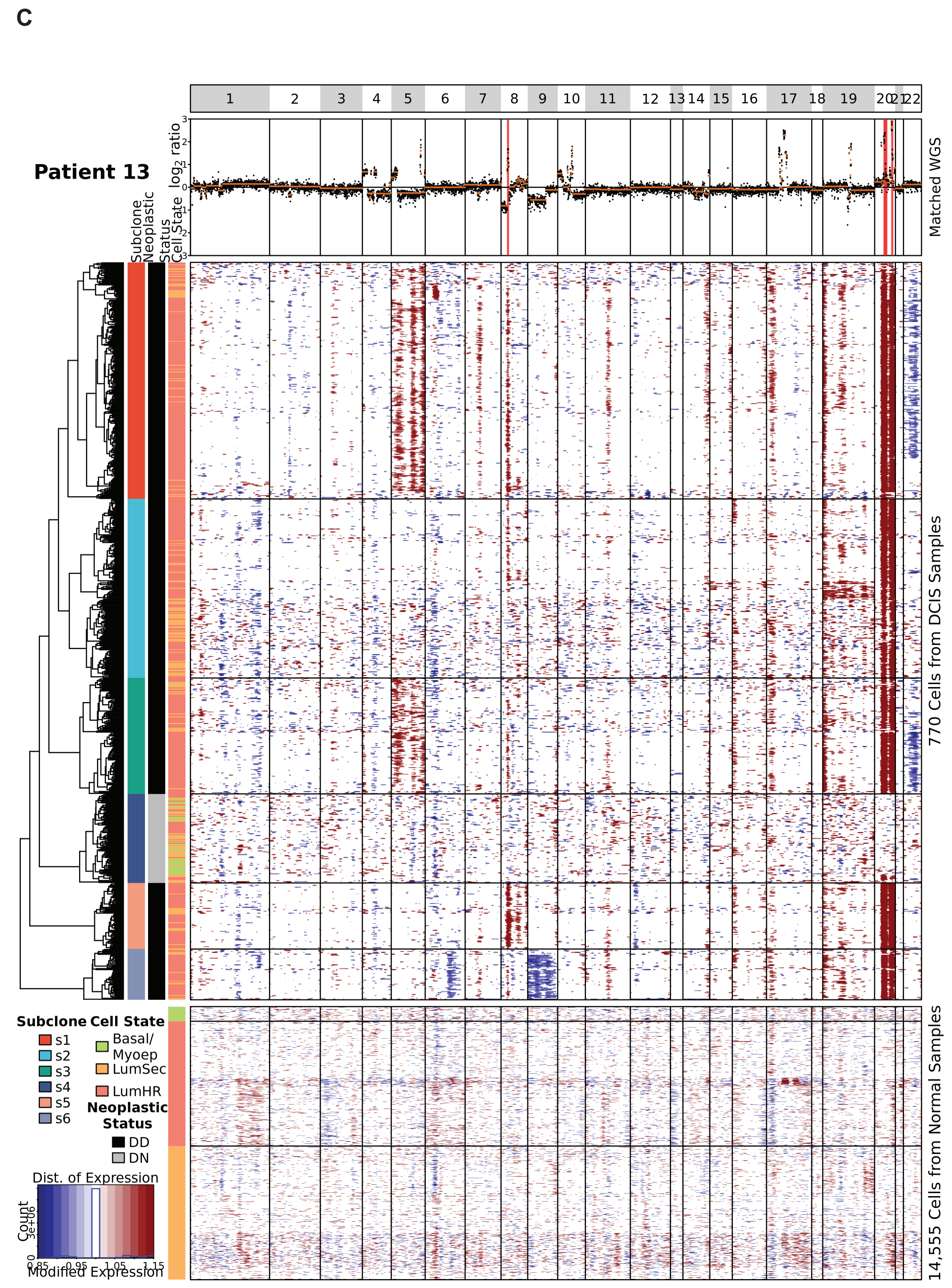

Supplementary Figure 2 (Cont.)

C

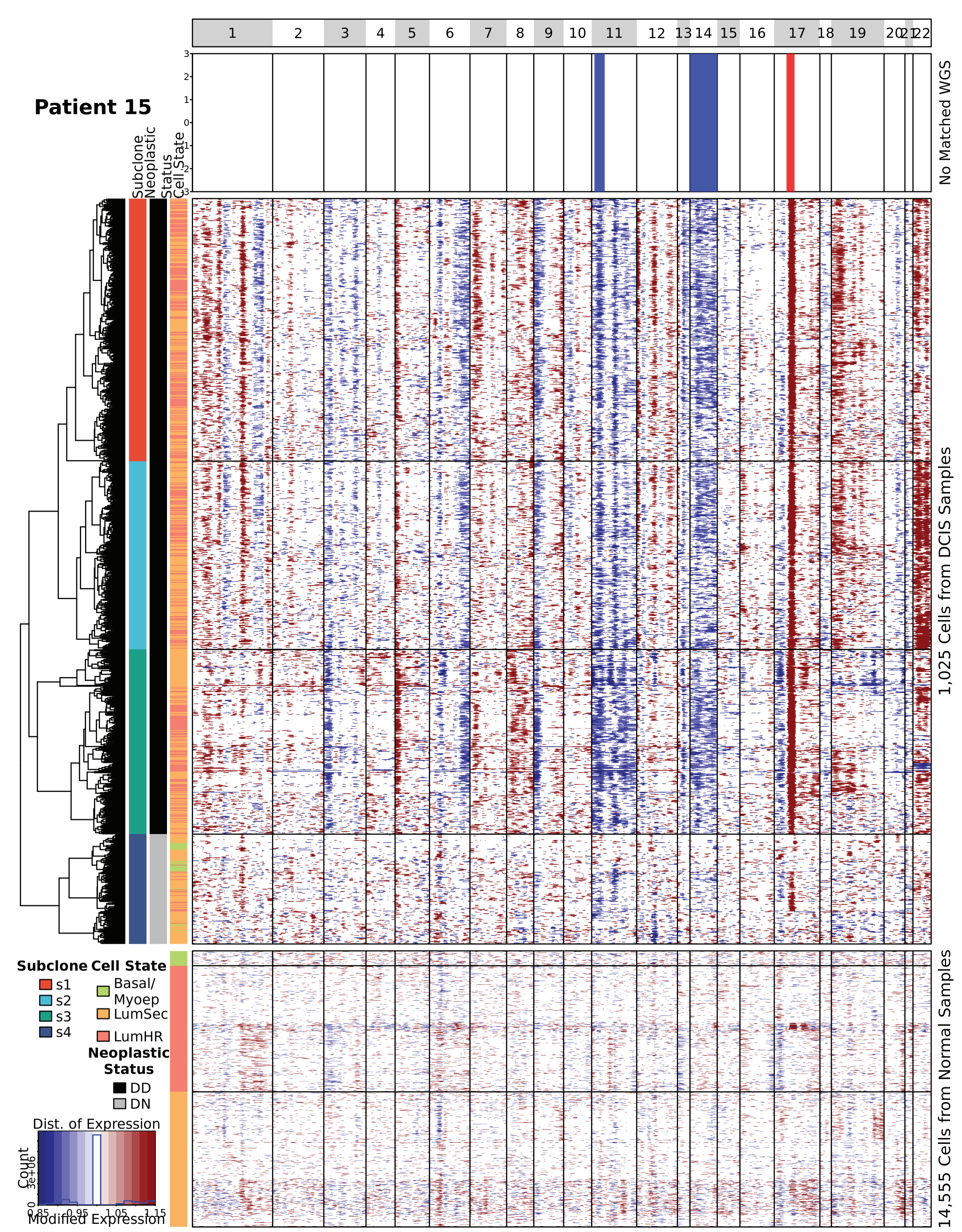

#### Supplementary Figure 2 (Cont.)

### C

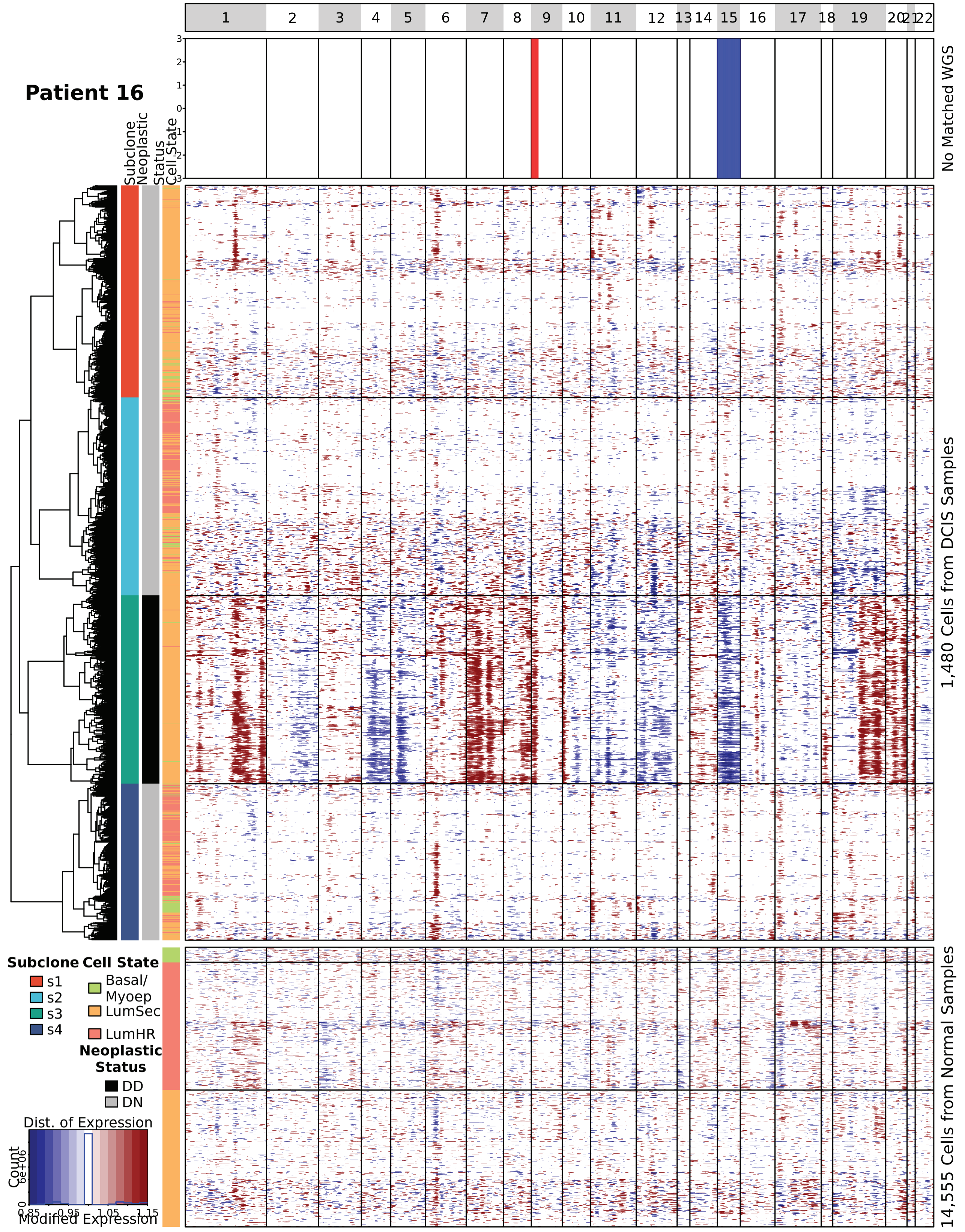

### Supplementary Figure 2 (Cont.)

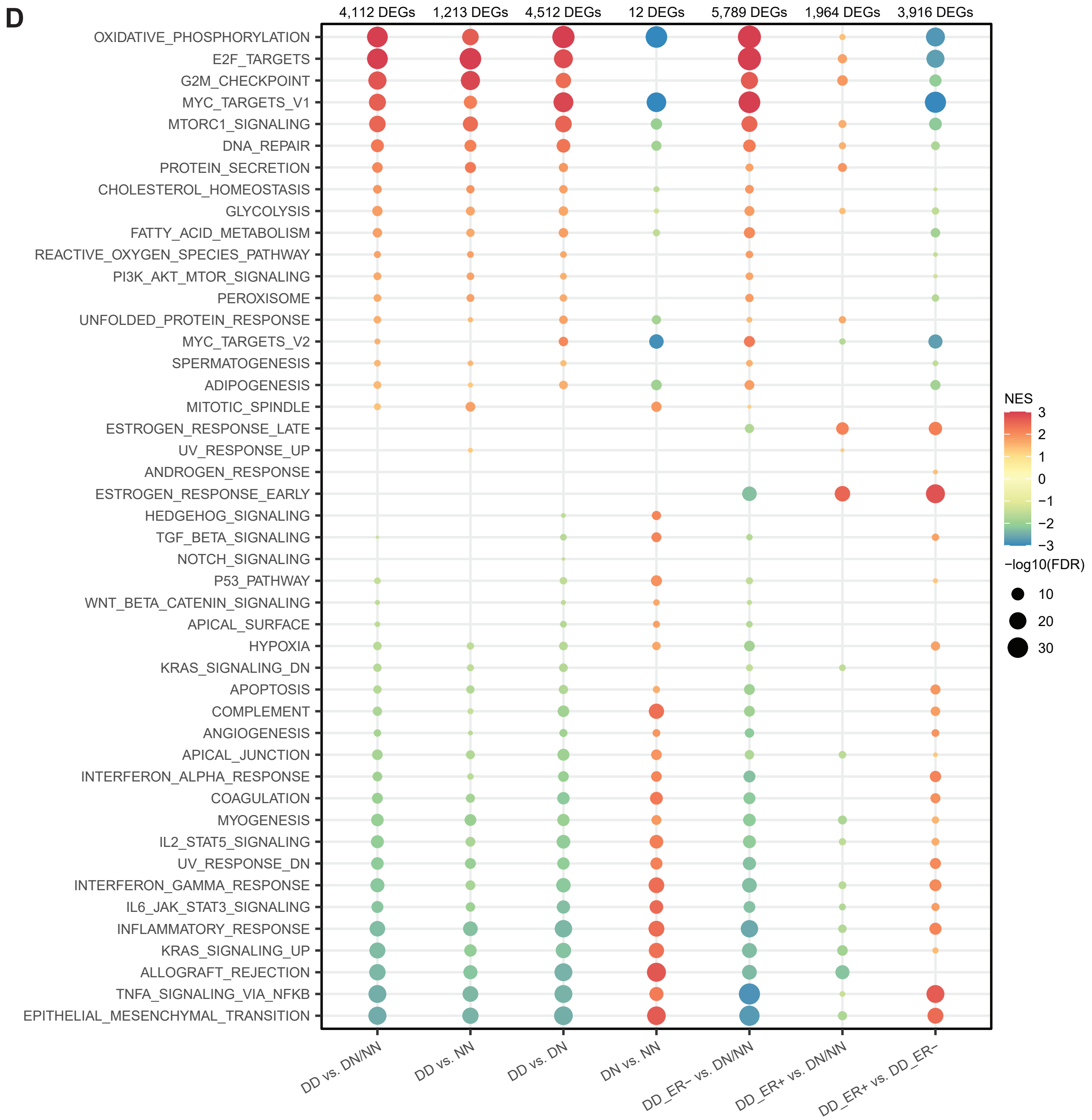


**Supplementary Figure 2**
**A)** Distribution of 11 mammary epithelial cell substates in each sample, with text in each cell indicating the raw cell count at the top and the cell count normalized by library size in parentheses. **B)** Histology of specimens used for scRNA-seq. Scanned hematoxylin and eosin stained section from the bisected facing half of the specimen dispersed for single cell analysis. Sections were scanned at 20X magnification and these screenshots from the scans show each entire specimen. **C)** Heatmaps of single cells with copy number status at each genomic locus, inferred from a low-pass WGS (upper panel) and inferred from single-cell DCIS (middle panel) libraries. A panel of 7 normal libraries were used as the reference (bottom panel). Each page shows an individual DCIS specimen. **D)** GSEA of all 50 Hallmark pathways between pseudo-bulk samples from neoplastic and non-neoplastic epithelial cells from breast tissue samples. The numbers of significant genes are indicated on top of the heatmap of the different comparisons. The color of the dots indicates the normalized enrichment score, and the size indicates significance with the negative log10 adjusted p-value.

### Supplementary Figure 3

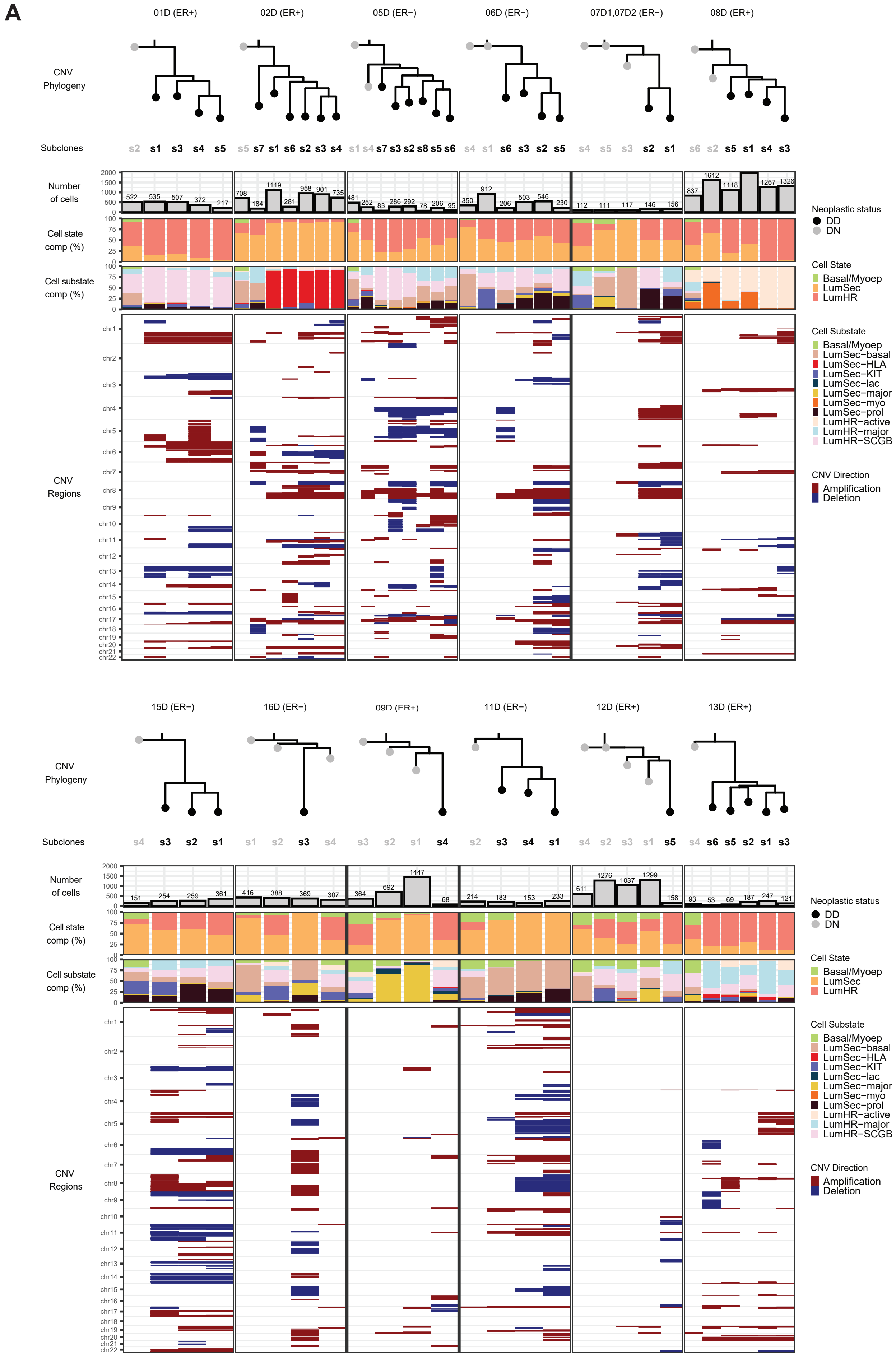

### Supplementary Figure 3 (Cont.)

**B**

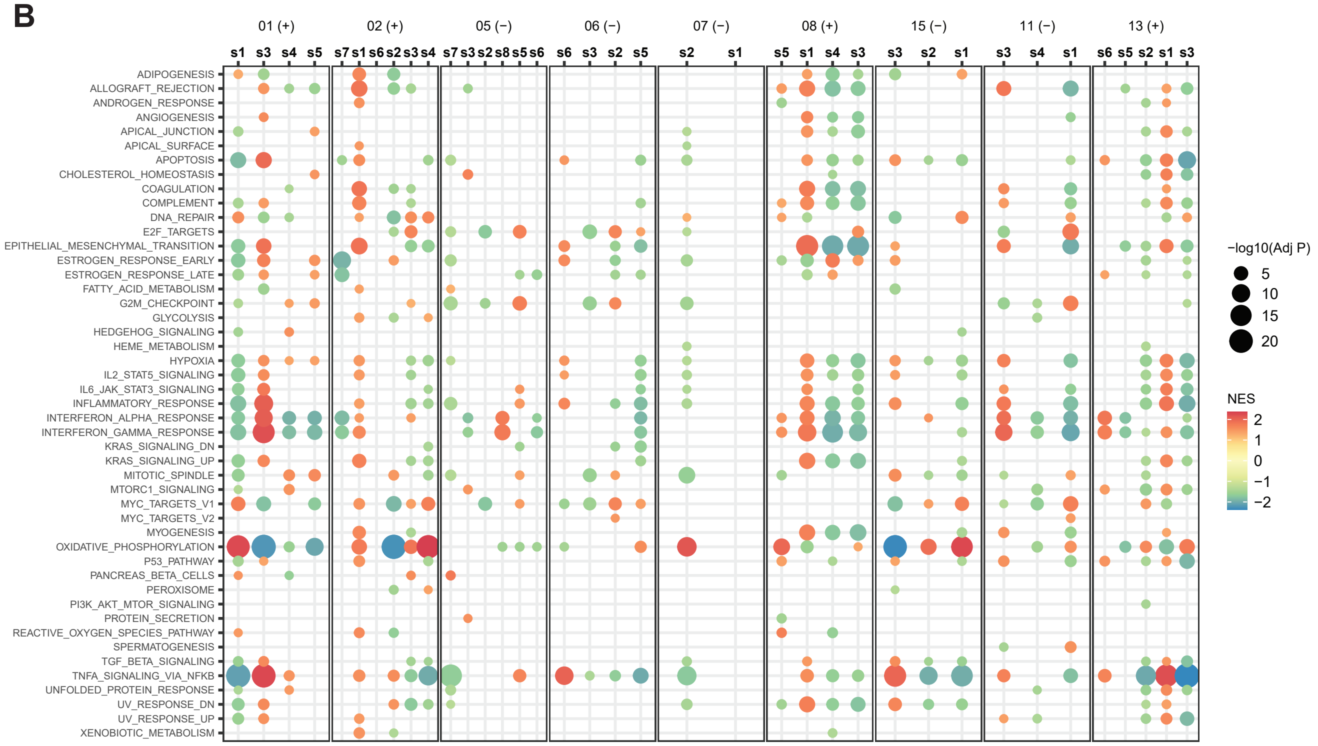

#### Supplementary Figure 3 (Cont.)

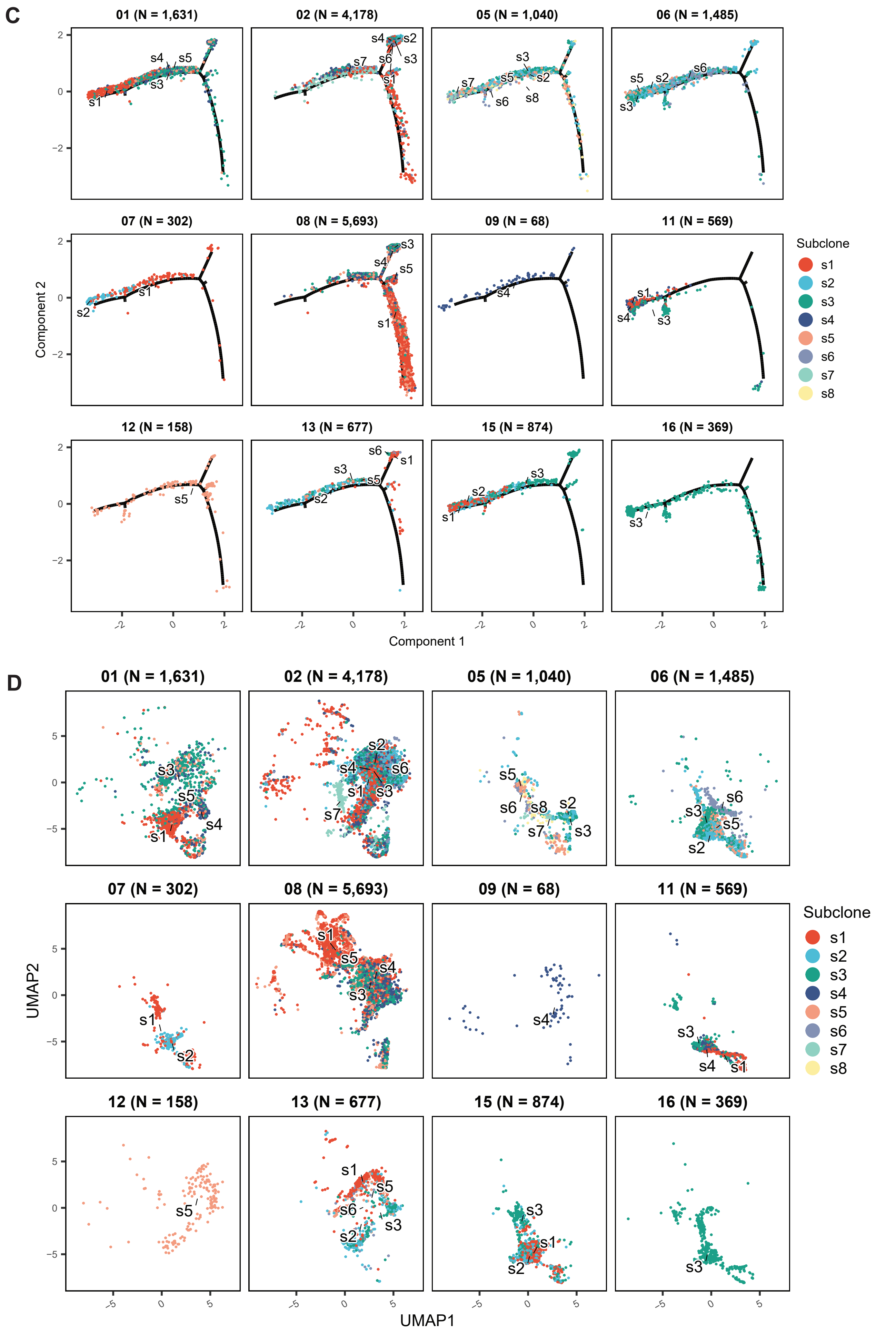

#### Supplementary Figure 3 (Cont.)

**E**

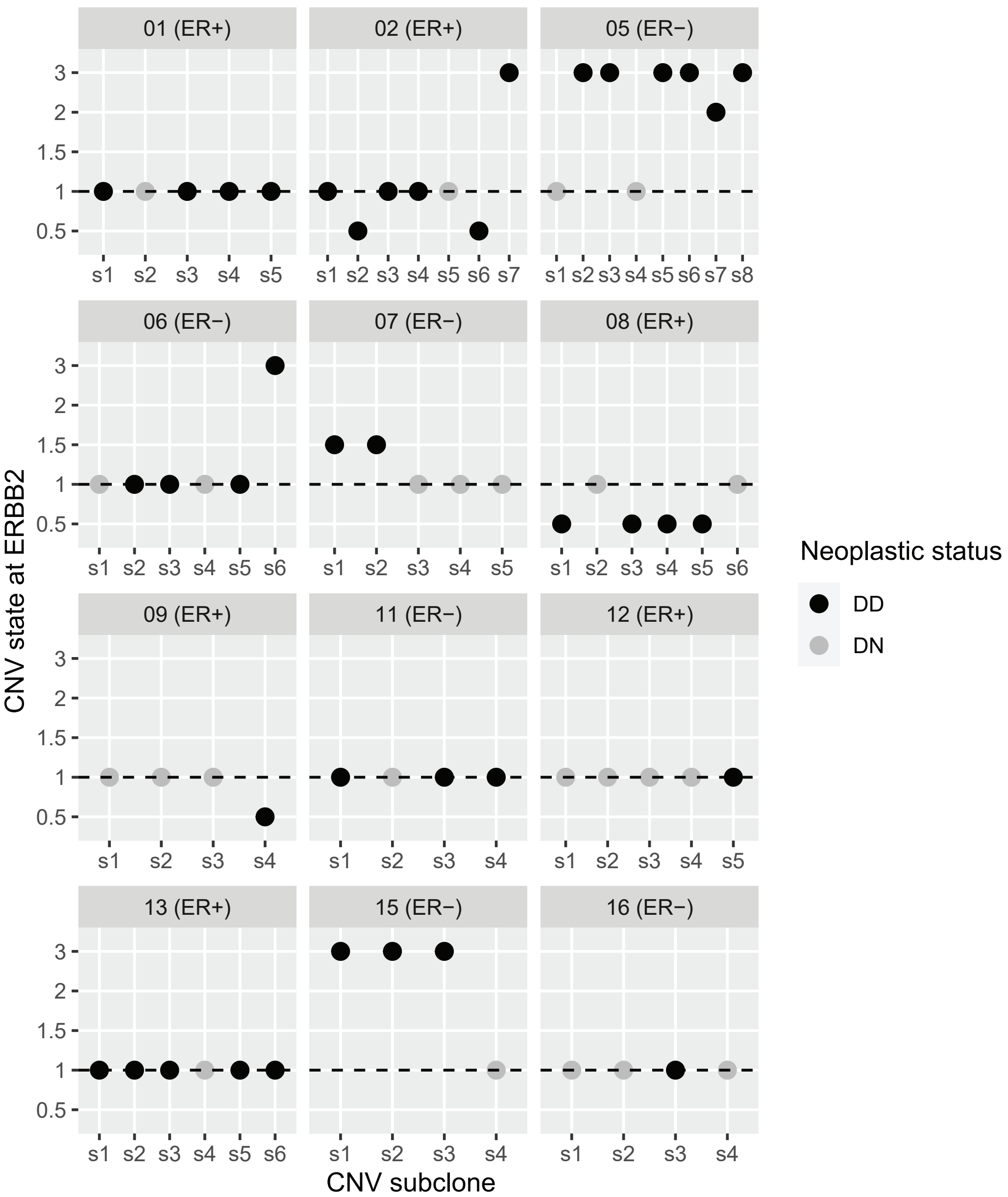


**Supplementary Figure 3**
**A)** Heterogeneity of CNV profiles in DCIS samples from each patient. For each of 12 patients, we show the subclone CNV phylogenetic trees with neoplastic status classifications as shown by tree tip and subclone label colors. The number of cells, cell state/substate compositions, and inferred CNV regions by subclone are shown in the bar charts and tile plots. **B)** GSEA results for a given DD subclone as compared to all the other DDs in the same patient DCIS tissue. Of the 12 patients, 9 had more than one DD subclone which was necessary to perform this analysis and generate the results shown here. **C)** and **D)** Position of DCIS genetic subclones in **C)** Pseudotime trajectory space and **D)** UMAP space. Subclones are colored separately for each case and reference to the phylogenies indicate the identity of the subclones. The pseudotime trajectory (**Figure 3I**) and UMAP (**Figure 3A**) were constructed from all scRNA-seq epithelial cells. **E)** Inferred CNV state at ERBB2 in DCIS subclones from each patient. Subclone neoplastic status is shown by point color and the patient's ER status is indicated in the panel headers.

### Supplementary Figure 4

**A**

| | Cell State | Cell Substate | Markers |
|---|---|---|---|
| 1 | B cells | B cells | IGKC,IGLC2,IGHA1,IGLC3,JCHAIN,IGHA2,IGHG1,IGHM,IGHG3,IGLL5,IGHG2,IGHG4,MZB1,IGHD,SSR4,CD79A,CD37,DERL3,HLA−DRA,HERPUD1 |
| 2 | Basal/Myoep | Basal/Myoep | TAGLN,KRT14,ACTA2,TPM2,KRT17,MT2A,DST,MYLK,MT1X,CXCL14,KRT5,APOE,SPARCL1,ACTG2,C2orf40,CALD1,CNN1,MT1E,MYL9,NNMT |
| 3 | Fibroblasts | Fibroblasts | DCN,APOD,CFD,TNFAIP6,LUM,COL1A2,COL1A1,COL3A1,MMP3,GSN,FBLN1,CCDC80,MEG3,SFRP2,COL6A2,IGFBP6,IGF1,C1S,COL6A3,C1R |
| 4 | LumHR | LumHR−active | CXCL13,DIO2,TFF3,MYBPC1,EREG,SLC26A3,PTHLH,FASN,MS4A7,TFF1,PLCG2,PNMT,CA2,C2CD4A,GSTM3,KCNMA1,NNMT,ANKRD30A,EFHD1,RGS1 |
| 5 | LumHR | LumHR−major | S100A6,TNFRSF11B,AC097059.1,FAM107B,ITGA2,C15orf48,SYTL4,TUBA1C,IER5,NFKBIA,RBP1,SYTL2,KRT8,ATP1B1,SPINK1,PAWR,HMGA1,ARID5B,TNFRSF12A,S100A14 |
| 6 | LumHR | LumHR−SCGB | SCGB2A2,PIP,SCGB1D2,MUCL1,SCGB3A1,SERPINA1,TFPI2,APOD,CP,KRT23,HMGB3,SAA1,MGP,CA2,TAT,SOD2,CITED1,CYP4X1,TNFSF10,CCL2 |
| 7 | LumSec | LumSec−basal | PTN,WFDC2,MGP,RARRES1,SCGB2A2,KRT6B,CXCL8,S100A8,CCL2,S100A6,S100A9,KRT14,KRT17,KRT81,S100A10,LCN2,PI3,KRT23,ADIRF,LTF |
| 8 | LumSec | LumSec−HLA | CCL20,PI3,SAT1,CD74,TMSB4X,HSPB1,RBP1,SLPI,CRABP2,IER3,HIST1H1C,SPARCL1,HLA−DRB1,NFKBIA,SELENOM,HLA−DRA,SNCG,IFI27,SERPINB1,ANXA1 |
| 9 | LumSec | LumSec−KIT | PLCG2,MAFB,SOX4,JUN,HES1,HEXIM1,CSKMT,FOSB,HIST1H4C,GOLGA8A,FOS,SLC38A2,IER5L,AL355075.4,HSPA2,NR4A1,HSPA1B,XIST,IGKC,AC023157.3 |
| 10 | LumSec | LumSec−lac | LALBA,CSN1S1,CSN2,CSN3,LYZ,SPP1,IGKC,LTF,FDCSP,IGHA1,FABP3,ZG16B,CLU,PLIN2,XDH,S100A1,PIGR,HLA−DRA,CEL,NUPR1 |
| 11 | LumSec | LumSec−major | SCGB3A1,CYP24A1,MSMO1,MMP7,TM4SF1,HMGCS1,PDCD4,CALML5,JUND,RPS18,RACK1,IDI1,RPLP0,RPS21,FAM3B,DBI,PIGR,RPS8,RPS28,MT−ND6 |
| 12 | LumSec | LumSec−myo | ACTA2,TAGLN,DST,TPM2,MT2A,MYLK,KRT14,CXCL14,KRT17,APOE,SPARCL1,C2orf40,CNN1,ACTG2,MYL9,MT1E,NNMT,IGFBP2,THBS1,POSTN |
| 13 | LumSec | LumSec−prol | HMGB2,TUBA1B,HMGN2,TOP2A,PTTG1,UBE2C,HIST1H4C,TYMS,NUSAP1,PCLAF,HMGB1,PCNA,TUBB,IGKC,H2AFZ,DUT,CDK1,CENPW,TK1 |
| 14 | Lymphatic | Lymphatic | CCL21,TFF3,MMRN1,CAVIN2,CLDN5,LYVE1,TFPI,PPFIBP1,GNG11,ECSCR,ANGPT2,PROX1,CD9,FABP4,FABP5,AKAP12,RAMP2,CAV1,EFEMP1,S100A10 |
| 15 | Myeloid | Myeloid | HLA−DRA,IL1B,HLA−DPA1,HLA−DPB1,HLA−DRB1,CD74,CCL3,HLA−DQA1,C1QB,C1QA,RNASE1,FCER1G,TYROBP,LYZ,CCL4,HLA−DQB1,GPR183,CTSB,CD163,C1QC |
| 16 | Perivascular | Perivascular | RGS5,C11orf96,MT1A,IGFBP5,STEAP4,MYL9,IGFBP7,ADIRF,ADAMTS4,TAGLN,PDK4,NDUFA4L2,GADD45B,RGS16,ADAMTS1,MYH11,NR2F2,CCL2,NOTCH3,MCAM |
| 17 | T cells | T cells | IL7R,CCL5,PTPRC,CXCR4,GNLY,CD2,SRGN,NKG7,KLRB1,ARHGDIB,CD3D,CREM,TRBC2,LEPROTL1,CD7,SARAF,CD52,SYTL3,CNOT6L,CST7 |
| 18 | Vascular | Vascular | SELE,ACKR1,FABP4,STC1,CLDN5,ANGPT2,CSF3,IFI27,ADGRL4,ADAMTS9,AQP1,C2CD4B,GNG11,TM4SF1,SPARCL1,CD93,VWF,RBP7,PECAM1,SPRY1 |

**B**

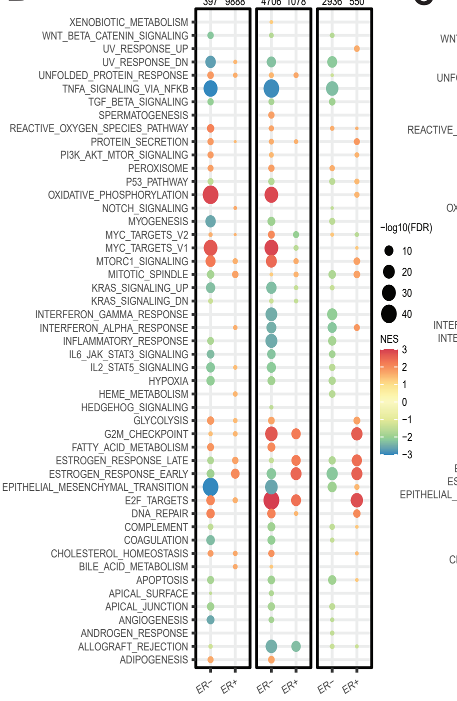


**C**

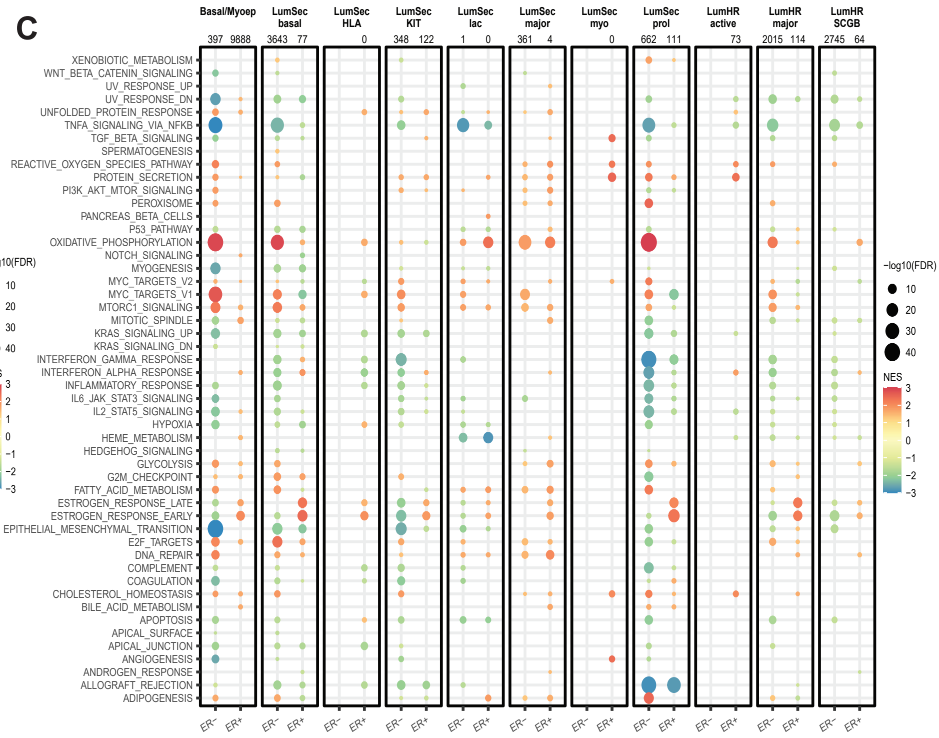


**Supplementary Figure 4. Mammary cell state and substate analysis.**
**A)** Genes used to define cell types and mammary epithelial cell states and substates. **B)** GSEA of all 50 Hallmark pathways between pseudo-bulk samples from DN/NN and DD of three major cell states in ER- DCIS and ER+ DCIS, analyzed separately. **C)** GSEA analysis as in **B)** but for the 11 mammary substates. The color of the dots indicates the normalized enrichment score, and the size indicates significance with the negative log10 adjusted p-value.

### Supplementary Figure 5

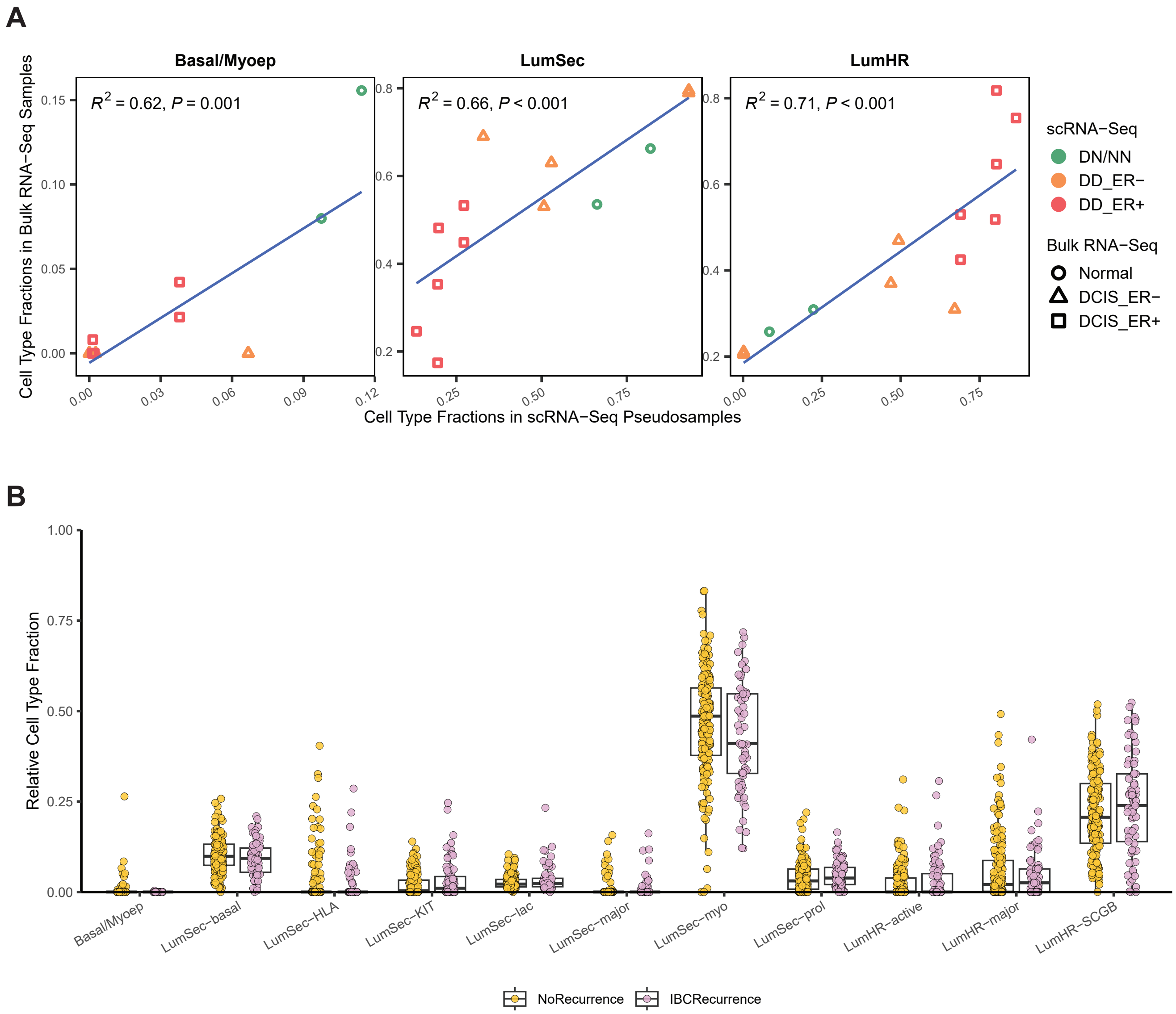


**Supplementary Figure 5. Bulk sample cell fraction analysis.**
**A)** Correlation between relative cell fractions from the 13 pseudo-bulk samples (5 DD ER-, 6 DD ER+, and 2 DNNN) derived from the 12 scRNA-seq specimens (10 DCIS, 2 Normal) and imputed relative cell fractions from 13 patient-matched FFPE bulk RNA-seq samples (5 ER- DCIS, 6 ER+ DCIS, and 2 normal), across 3 major epithelial cell states (Sample set 1). **B)** Analysis of imputed cell fractions based on 11 epithelial substates for 163 DCIS that did not progress or recur and 69 DCIS that progressed to IBC from Sample set 3.

#### Supplementary Figure 6

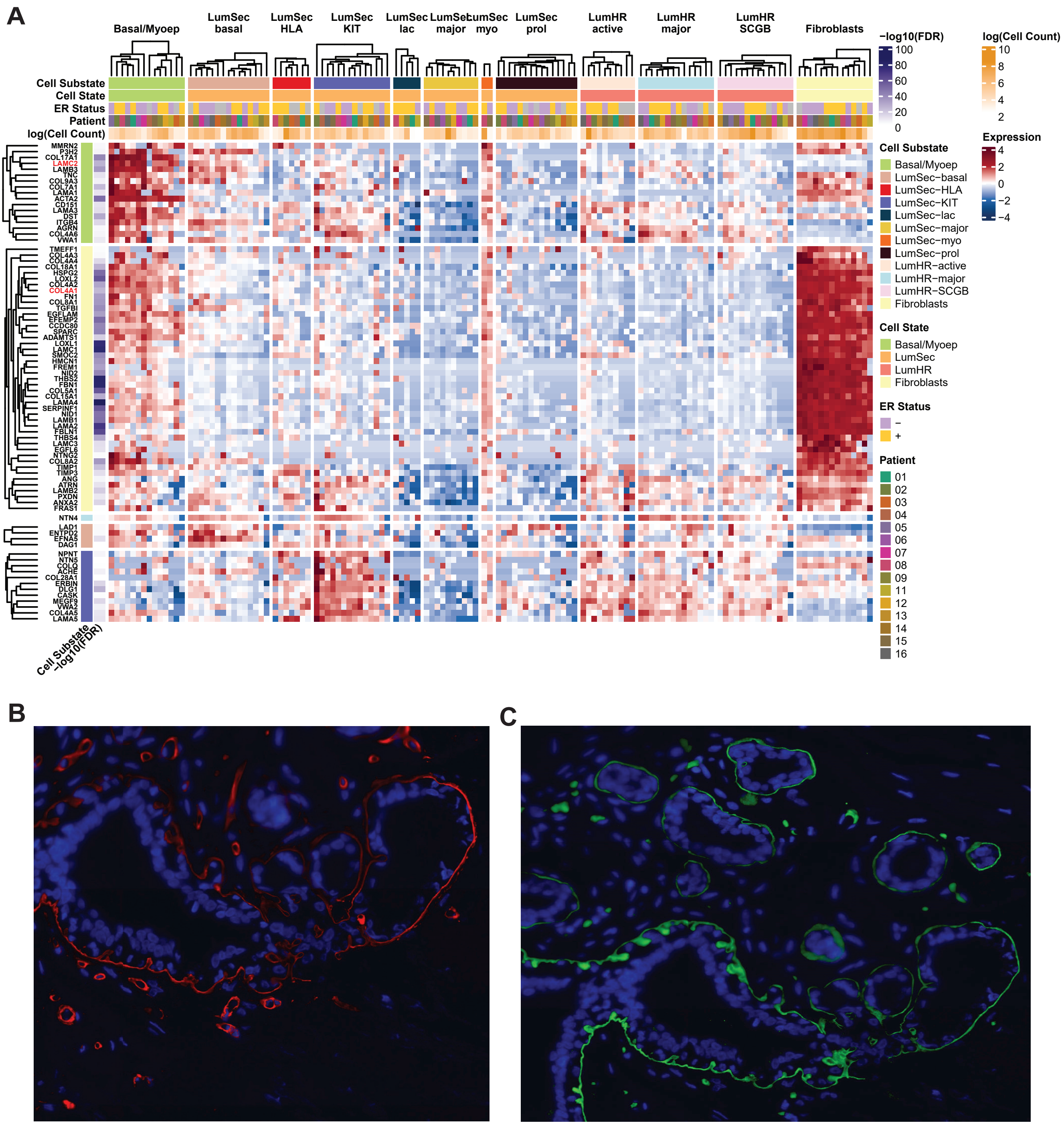


**Supplementary Figure 6. Expression of basement membrane genes.**
**A)** Relative expression of BM genes across 11 epithelial cell substates and fibroblasts within each pseudo-bulk sample from scRNA-seq data (Sample set 1). Displayed genes were most significantly up-regulated (adj. $p < 0.01$) in each specific cell type compared to all other cell types. **B)** Immunofluorescent staining of COL4A1 (red) in normal breast and **C)** LAMC2 (green) in normal breast with blue DAPI nuclear stain (Sample set 4).
